# Invisible shield: Sprayable supramolecular antimicrobial microscale films for preventing wound and medical device infections

**DOI:** 10.64898/2026.04.10.717441

**Authors:** Yijie Li, Skander Hathroubi, Ophélie Heck, Lucie Lieu, Lauriane Petit, Xavier Wurtz, Abdessalem Rekiki, Aurore Gaudin, Naomi Canourgues, Derry Mercer, Merve Tunali, Bernd Nowack, Philipp Meier, Giacomo Reina, Peter Wick, Marjan Safarzadeh, Altan Demircan, David Grossin, Christophe Drouet, Thibaut Soubrié, Tetiana Holdanova, Mélanie Kremer, Noémie Willem, Sarah Jester, Aurélia Cès, Cynthia Calligaro, Baptiste Letellier, Agnes Dupret-Bories, Philippe Lavalle, Nihal Vrana Engin

## Abstract

Wound and device-associated infections remain difficult to eradicate because biofilms block host immunity and antibiotics, accelerating chronicity and resistance. Here, we present a portable, low-cost dual-syringe spray that deposits an ultra-thin, self-assembling antimicrobial film directly on wounds and implant surfaces. The device co-delivers oppositely charged hyaluronic acid (HA) and a cationic antimicrobial peptide (polyarginine, PAR30), which rapidly form a conformal nanometric polyelectrolyte complex at the tissue-material interface. Molecular dynamics simulation revealed pronounced positional heterogeneity within the PAR30/HA complex and identified an N-terminal arginine as a dominant interaction hotspot. The resulting coating adheres to diverse substrates, kills bacteria on contact, prevents biofilm formation, and sustains antimicrobial efficacy. *Across vitro* assays and murine wound infection models, treatment produced 4 to 5 log reductions in bacterial burden against methicillin-resistant *Staphylococcus aureus* and Gram-negative pathogens, including *Pseudomonas aeruginosa* and *Escherichia coli*. The formulation is biocompatible, did not increase cutaneous inflammation or IL-6 levels *in vivo*, and reduced post-surgical pain and motor deficits in a mouse incision model. To our knowledge, this is the first antimicrobial treatment system applicable to both tissues and medical devices. Developed under a safe-and-sustainable-by-design approach, this technology combines biocompatible components, nanometric coating for minimal material use, and a simple syringe-based delivery device, offering a scalable, antibiotic-free strategy for wound care and medical device infection prevention.

**Graphical abstract:** 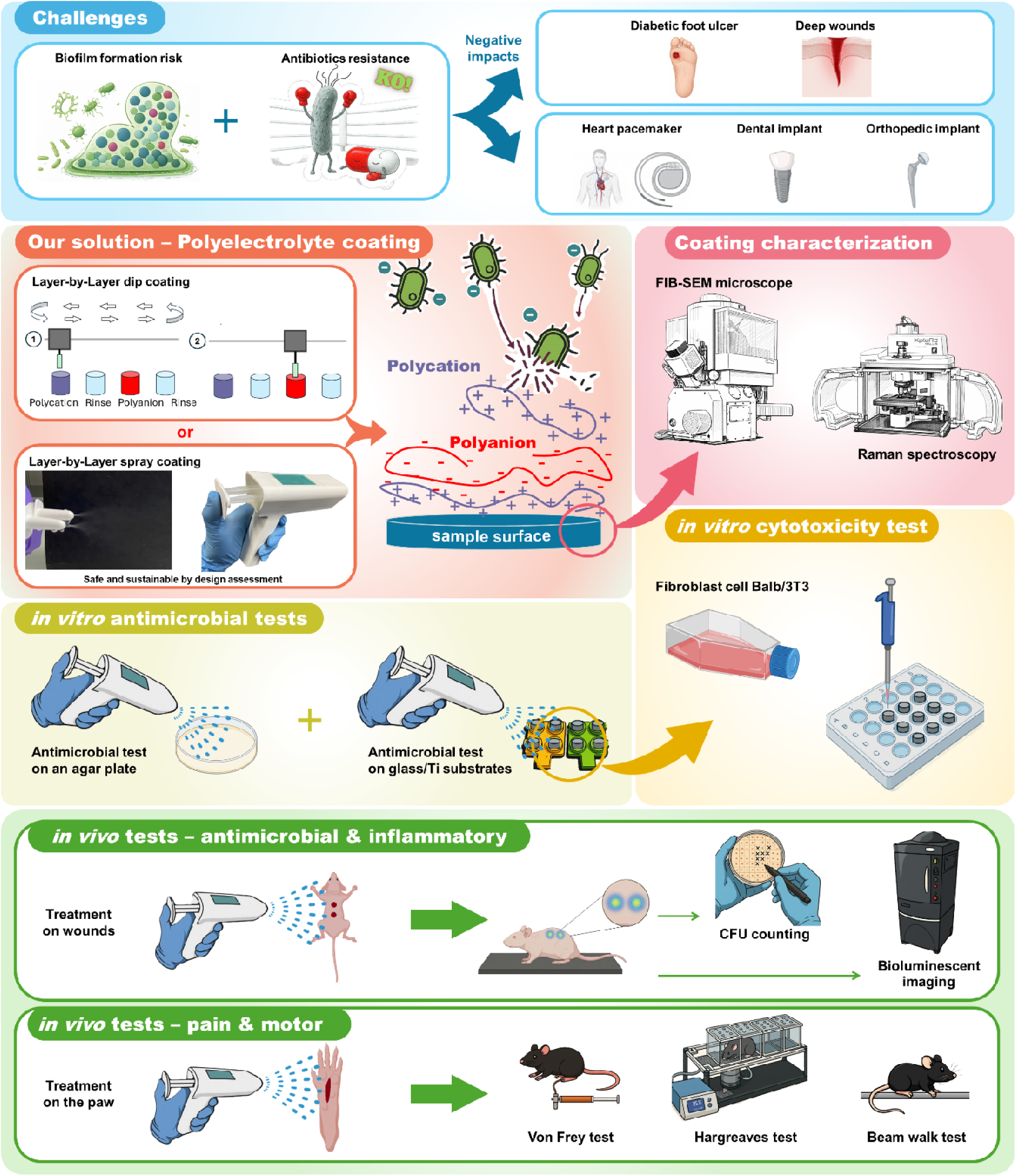

## Introduction

Chronic wounds represent a significant global health concern with major patient-related and economic consequences ^1^. Such wounds fail to progress through the normal stages of healing within an appropriate timeframe, typically considered to be three months, or fail to achieve permanent anatomical and functional integrity upon closure ^2^. The terminology surrounding these wounds is inconsistent, with terms such as "non-healing wounds" and "ulcers" often used interchangeably. These full-thickness wounds exhibit slow healing and are frequently complicated by underlying comorbidities. The prevalence of chronic wounds is notable, with an estimated 1–2% of individuals in developed countries likely to develop such wounds during their lifetime, a rate expected to rise with the aging population due to the decline in wound healing associated with age. Chronic wounds contribute to significant disability, creating a vicious cycle where impaired mobility exacerbates the risk of wound recurrence and severity.

Complications associated with chronic wounds are severe and can include infections such as cellulitis, infectious phlebitis, necrosis, hemorrhages, and, in extreme cases, lower-limb amputations ^3^. Diabetic foot ulcers (DFUs), a common type of chronic wound, illustrate the severity of this problem, with a 40% infection rate among diabetic patients, leading to a high risk of gangrene and amputation. The global burden of diabetes, a major risk factor for chronic wounds, is escalating at an alarming rate: in 2021, approximately 537 million people were living with diabetes, a figure projected to rise to 643 million by 2030 and 783 million by 2045 ^4^ .

The challenge of treating chronic wounds is compounded by the presence of pathogenic bacteria, which colonize the wound surface. Infections often involve a variety of bacterial species, including *Staphylococcus* spp., *Streptococcus* spp., *Pseudomonas aeruginosa*, and other Proteobacteria in the form of small to large aggregates and individual cells ^5,6^. These microorganisms can exist in planktonic forms or as biofilms, the latter presenting a major challenge due to their physical protection from the host immune response and their increased tolerance to antibiotics ^7^. Biofilms are complex bacterial communities embedded within an extracellular polymeric substance (EPS) matrix, providing a protective environment that significantly enhances bacterial survival and locally modifies the wound microenvironment. The transition from planktonic to biofilm states dramatically increases antibiotic tolerance, up to 10,000-fold compared to free-floating bacteria, due not only to the physical barrier provided by the EPS matrix, but also to multiple physiological and genetic factors ^8^. Biofilm cells often exhibit reduced metabolic activity, entering slow-growing or even dormant states where many antibiotics, which typically target active processes, like cell wall synthesis or protein production, are less effective. The high cell density and proximity of bacteria promote the spread of antibiotic resistance genes, thus contributing to the emergence of multidrug-resistant (MDR) strains and persistent infections ^9^. In chronic wounds, the presence of biofilms leads to delayed healing, recurrent infections, and severe clinical challenges, further complicated by the nutrient-rich environment created by wound exudates ^7,10^.

Surgical site infections (SSIs) represent a significant complication in procedures involving indwelling medical devices, particularly due to the frequent formation of biofilms on implanted surfaces ^11^. These biofilms provide a protective environment for microorganisms, enhancing their resistance to both host immune responses and antimicrobial therapies; thereby making infections extremely difficult to eradicate. Consequently, device-related infections often necessitate removal and replacement of the implant, which significantly increases patient morbidity, mortality, and healthcare costs. This concern is particularly obvious in the context of cardiovascular implantable electronic devices (CIEDs), where the risk of adverse events (AEs) remains substantial. In a 2015 analysis, a cohort of 60,296 patients underwent invasive CIED procedures in Germany ^12^ . Within three months of the index procedure, 1,595 patients developed major CIED infections. Of these, 1,129 cases (1.87%) were associated with the generator pocket, while 466 cases (0.8%) involved the transvenous portions of the leads. The associated mortality rates were 8.4% and 15.24%, respectively ^12^. Similarly, a study in Canada has shown that 1.23% of CIED procedures resulted in SSIs^13^. Both studies underscore the significant increase in costs and clinical burden of CIED infections to the healthcare system ^12,13^.

Despite current advancements in wound care and therapeutic options, there are no solutions that can simultaneously treat the wounds (whether chronic or surgical-site wounds) and related medical devices (such as wound dressings, hernia meshes, or implantable devices). Removal of the implant is often necessary to control the infection, although the decision is made on an individual basis, depending on the morbidity of removal and the risk of reoperation ^14^. Thus, there remains an urgent need for versatile strategies to prevent infection and promote healing in both wounds and implantable devices. Given the crucial role of bacterial colonization and subsequent biofilm formation in chronic wound pathogenesis and implant failure, effective wound management and implantation procedures must prioritize the early prevention of bacterial adhesion and biofilm development.

Previously, we have developed antimicrobial polyelectrolyte coatings that can be constructed on a wide range of substrates by a Layer-by-Layer (LbL) dip coating method ^15,16^. The conventional LbL dip-coating process involves immersion of substrates into polycation and polyanion solutions alternately, with rinsing steps in between. Automated LbL dipping robots have been developed to ensure reproducible, high-quality coatings ^17,18^. Recently, we have demonstrated the efficacy of such coatings on the protection of hernia meshes^15^ and wound dressings *in vitro* and *in vivo* ^19^. However, this approach can only fulfil the needs of substrates such as implantable medical devices under controlled conditions and cannot address potential contamination in surgical or clinical environments, nor enable direct application on the wound. This limitation highlights the need for *in situ* coating application systems.

Our research addresses this critical need through the development of a portable antimicrobial spray system capable of applying coatings to a wide range of substrates and surfaces (e.g. living tissues, implantable medical devices, and surgical tools) and in diverse environments (e.g. the clinic, operating room, and household settings for wound management). Given that products in the healthcare sector can adversely affect both the environment and human health ^20^, sustainability considerations are central to the proposed solution. In this context, the European Commission introduced the Safe-and-Sustainable-by-Design (SSbD) Framework in 2022 to support the sustainable innovation and development of chemicals and materials ^21^, with a revised version released in 2025 ^21^. Accordingly, a simplified SSbD assessment was conducted during the early phase of development on the solution to support its sustainable innovation. The main aim of the initial SSbD assessment is to identify potential hotspots for safety and sustainability issues beyond current regulations, to assess the initial safety and sustainability performance, and to define the steps that require improvement or possible advancements. The sprayable formulation is designed to inhibit initial bacterial attachment, reducing bacterial load to promote wound healing and implant integration by providing a biocompatible, non-inflammatory inducing, and protective layer. This coating formulation incorporates biodegradable polymers ^15,22,23^, offering several advantages for wound care applications, such as antimicrobial, biocompatible, lack of induced inflammation or pain, and support of wound hydration. One of the main advantages of the sprayable coating is that it can be easily applied to irregular surfaces, forming ultra-thin, transparent films that maintain a moist environment supportive of tissue regeneration while simultaneously serving as reservoirs for antimicrobial agents to prevent bacterial proliferation. Specifically, the coating is formulated with antimicrobial peptides (AMPs) as polycations, which are highly effective as a bactericidal system without the development of resistance by bacteria^15,24^. Hyaluronic acid (HA) is also incorporated into the formulation as the polyanion, known for its beneficial properties in wound healing, inflammation reduction, and tissue regeneration ^23,25,26^ . The combination of AMPs and HA on the targeted substrates is expected to create a barrier that prevents bacterial attachment by targeting bacterial membranes through electrostatic interactions, regardless of bacterial physiological status.

The portable, easy-to-use spray system aims to enable polyelectrolyte coatings to be applied within seconds. This device is not only time-efficient but also user-friendly. Its portability should allow application on complex surfaces, including direct application on several substrates (for example, multi-material implantable medical devices) or onto biological tissues such as wounds. In this study, we introduce a newly developed portable spray device capable of simultaneously delivering polycation and polyanion solutions without requiring intermediate buffer rinsing steps. We evaluated the spray efficacy, antimicrobial efficacy, cytocompatibility, and biomedical material compatibility of the coatings produced by this device through both *in vitro* and *in vivo* experiments. Coatings were applied to various materials, including cover glass slides, medical-grade titanium, medical-grade silicone, medical textiles, and agar plates (used here as a mimic of biological substrates). The spray was also tested in murine wound infection models. Furthermore, cytotoxicity and animal behavioral responses were assessed to validate the safety and potential clinical relevance of the spray system for biomedical applications, with potential pain-alleviating features for wound management. The spray is designed for controlled clinical environments such as hospitals and wound care centers. It is intended to be administered by qualified healthcare professionals to ensure safety and coating efficacy for chronic wound management and to deliver antimicrobial coating onto implantable medical devices in a customized manner.

## Results

### LbL dip coating background

LbL dip coatings are constructed through the sequential adsorption of oppositely charged polyelectrolytes, driven by multiple electrostatic interactions. In the standard protocol, substrates are alternately immersed in polycation solutions and polyanion solutions with a buffer rinsing step between each deposition. This process can be carried out via a robotic dipping system (Figure 1.a, b). A typical 24-bilayer film, denoted as (PAR30/HA), and built from alternating polycations (PAR30) and polyanions (HA), requires approximately 6 to 13 hours to complete ^15,16^. Despite the time-intensive nature of this method, the resulting coatings are highly reproducible, stable, and bioactive against Gram-negative and Gram-positive bacteria ^15^.

**Figure 1.**
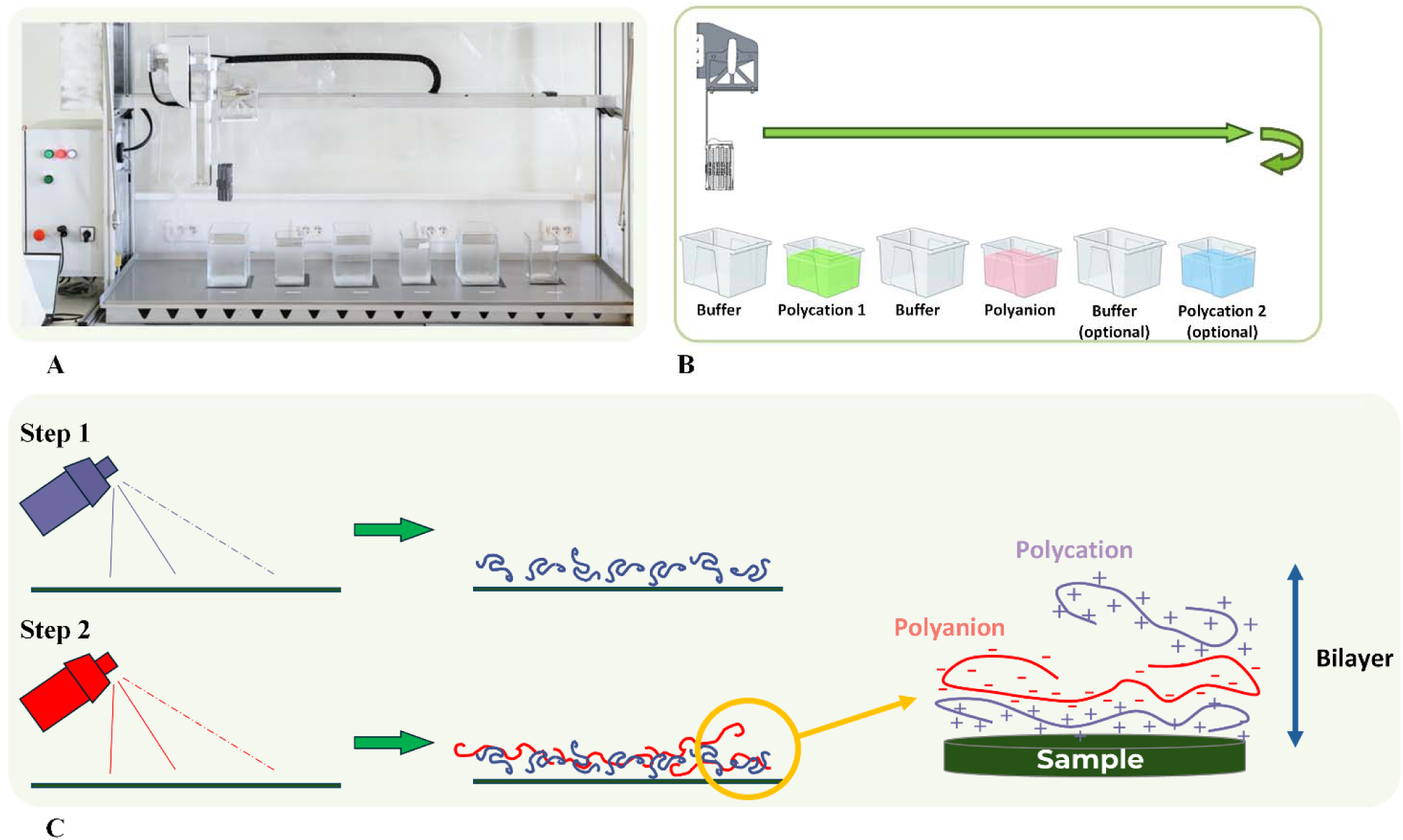
LbL dip coating process. (**A)**. Spartha Medical industrial standard dipping robot. (**B)**. Spartha Medical industrial dipping robot LbL coating process. (**C)**. One of the traditional spray coating methods.

Despite the high degree of architectural control, the duration, complexity, and reliance on robotic equipment associated with dipping limit its use at the point-of-care. An alternative approach uses separate spray bottles, spraying polycation and polyanion solutions respectively or simultaneously to fabricate the multilayer polyelectrolyte coatings on substrates ^27^ (Figure 1. C).

Although conventional LbL dipping yields reproducible and bioactive coatings, its sequential immersion and rinsing steps make the process too slow and operationally demanding for direct wound care and point-of-care use. Existing spray-based alternatives simplify deposition, but disconnected spray systems provide limited control over synchronized delivery and coating asymmetry. To address this, we developed a compact dual-container spray device that translates LbL deposition into seconds per layer without rinsing, while remaining compatible with clinical workflows and good manufacturing practice (GMP) standards. This no-touch approach also enables spraying onto the wounds, adapts to irregular wound geometries that are challenging to cover with prefabricated dressings^28^, minimizes contamination risk, and rapidly generates tailored antimicrobial multilayers for infection prevention ^28^.

### Spray device design and development

Because the spray system is intended for direct application of two biopolymer solutions onto wounds and medical-device surfaces, early translational considerations from the LbL to the spray method were integrated during development. We first addressed the central practical limitation of LbL assembly by designing a portable device that could separately store and simultaneously spray the two components to form the coating *in situ*. The design study addressed the ergonomic development of a portable device capable of delivering two biopolymer solutions. The device was designed for intuitive, instruction-free operation (Figure 2). A dual-nozzle configuration maintains separation of the two solutions until spraying, where they mix to form a film. This mechanism ensures fine mist delivery, which should reduce patient discomfort by lowering pressure on wounds. The device operates through mechanical actuation for improved reliability and simplified manufacturing. A design specification table (Supplementary Table 1) defined the key parameters, performance criteria, and validation methods guiding iterative development to ensure functionality, biocompatibility, and clinical relevance. The resulting compact spray system meets both functional and clinical requirements by enabling simultaneous yet separate spraying of the two biopolymer solutions, forming a bi-layered antimicrobial coating optimized for healthcare use (Figure 2).

**Figure 2.**
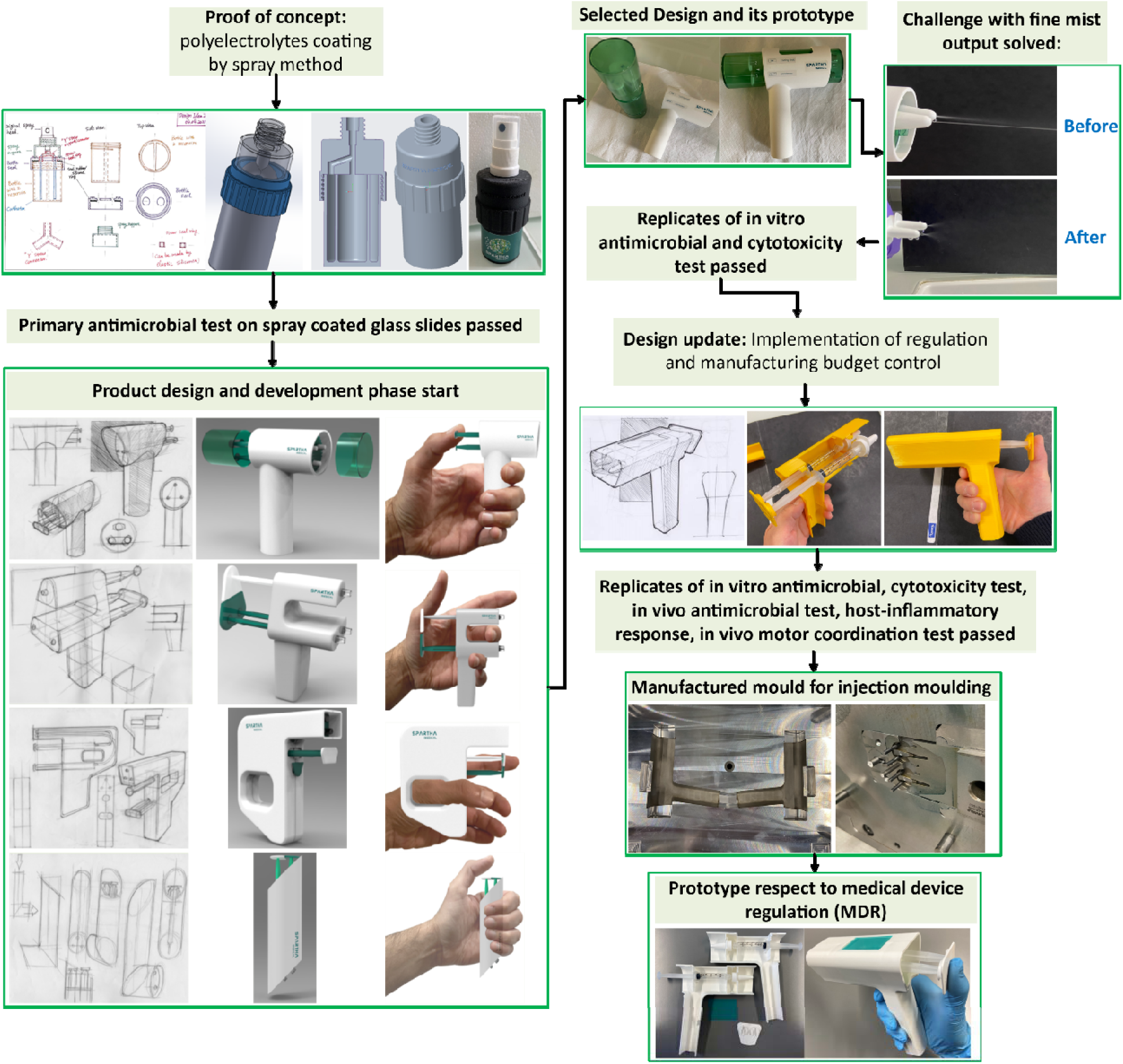
Schematic overview of the product design and development process. This illustration shows the antimicrobial polyelectrolyte spray coating applicator development process, from the iterative prototype design, early *in vitro* validation, to prototype optimization, regulatory implementation, and injection-molded manufacturing for a minimum viable product of a medical device.

The design and development steps were undergone in two phases: mechanism design and ergonomic optimization. Eco-design principals informed efforts to reduce material use and simplify manufacturing, while the McKinsey Design Index^29^ informed the use-centered ergonomic design process. Market analysis of existing fine-mist systems revealed that most applications rely on pump-based spray bottles. To validate the spray coating concept, a double-chamber bottle prototype using a commercial medical pump (AptarGroup, Crystal Lake, IL, USA) was modelled in SolidWorks and 3D-printed in Polylactic Acid (PLA) (Ultimaker S3) (Supplementary Figure S1. A). Usability tests unveiled some assembly complexity and inconsistent mist formation due to pressure loss. A second-generation concentric reservoir design was thus engineered, leading to a simplified and more compact assembly (Supplementary Figure S1. C). Preliminary *in vitro* antimicrobial testing confirmed activity (data not shown), albeit efficient coating required over 40 actuator presses, which undermined usability and costeffectiveness-. Consequently, the pump system was replaced with a syringe-based spray mechanism. A new ergonomic casement enabled simultaneous actuation of two syringes, in which each piston forces the solution through a micrometer-scale gap, releasing it into a fine, homogeneous mist (Supplementary Figure S1. D). Prototypes were 3D printed (Supplementary Figure S1. E, F), refined through iterative testing, and ultimately simplified by removing a replaceable chamber to ease manufacturing and reduce costs. Components were later injection-molded via computer numerical control (CNC) machined aluminum molds (Supplementary Figure S1 G, H) (Weil Industry, France). The design and development process of the applicator involves design, prototyping, and testing (Figure 2). A design specification table was developed during the prototyping process, the usability test, the *in vitro* test, and the *in vivo* test results contributed to the final decision on the primary product (Supplementary Table 1).

The final prototype integrates (i) two 2.5 mL medical-grade syringes (Terumo Corporation, Tokyo, Japan) as reservoirs, (ii) commercial nasal nozzles (AptarGroup, Crystal Lake, IL, USA) for atomization, and (iii) a custom nozzle-to-syringe adaptor enabling simultaneous discharge. The synchronized actuation allows lateral movement for spray-based LbL deposition without intermediate rinsing, simplifying traditional LbL processes while preserving sequential polymer layering.

With the view to allow actual manipulability and ease of use, the final system embodies an intentionally low-tech, yet clinically compliant system based on standardized medical components integrated into a compact ergonomic design. It is robust, cost-efficient, and suitable for rapid deployment in resource limited or emergency settings. By minimizing mechanical complexity, it ensures reliable handling, scalable production, and compliance with sterility requirements. This final iteration balances usability, technical feasibility, and antimicrobial performance, supporting future industrialization and field application of *in situ* biopolymer coatings.

### *In vitro* evaluation of antimicrobial efficacy using the novel spray device

Based on the preliminary testing from the double-chamber spray bottle (data not shown), a new functional spray device incorporating a syringe–nozzle system was developed and prototyped. This prototype was evaluated through *in vitro* antimicrobial testing using 10 mg/mL ε-poly-L-lysine (εPLL) and 1 mg/mL HA solutions. To establish user guidelines, different spray parameters were examined, including spray patterns (single-layer vs. three-layer application) and spray distances (10 cm vs 15 cm) (Figure 3). A spray distance between the spray nozzle and the substrates of 10 to 15 cm is clinically recommended to ensure safe and effective wound care, and to avoid complications like air embolism by allowing proper atomization and even coverage while minimizing tissue damage and patient discomfort, as supported by clinical guidelines and product instructions ^30^.

**Figure 3.**
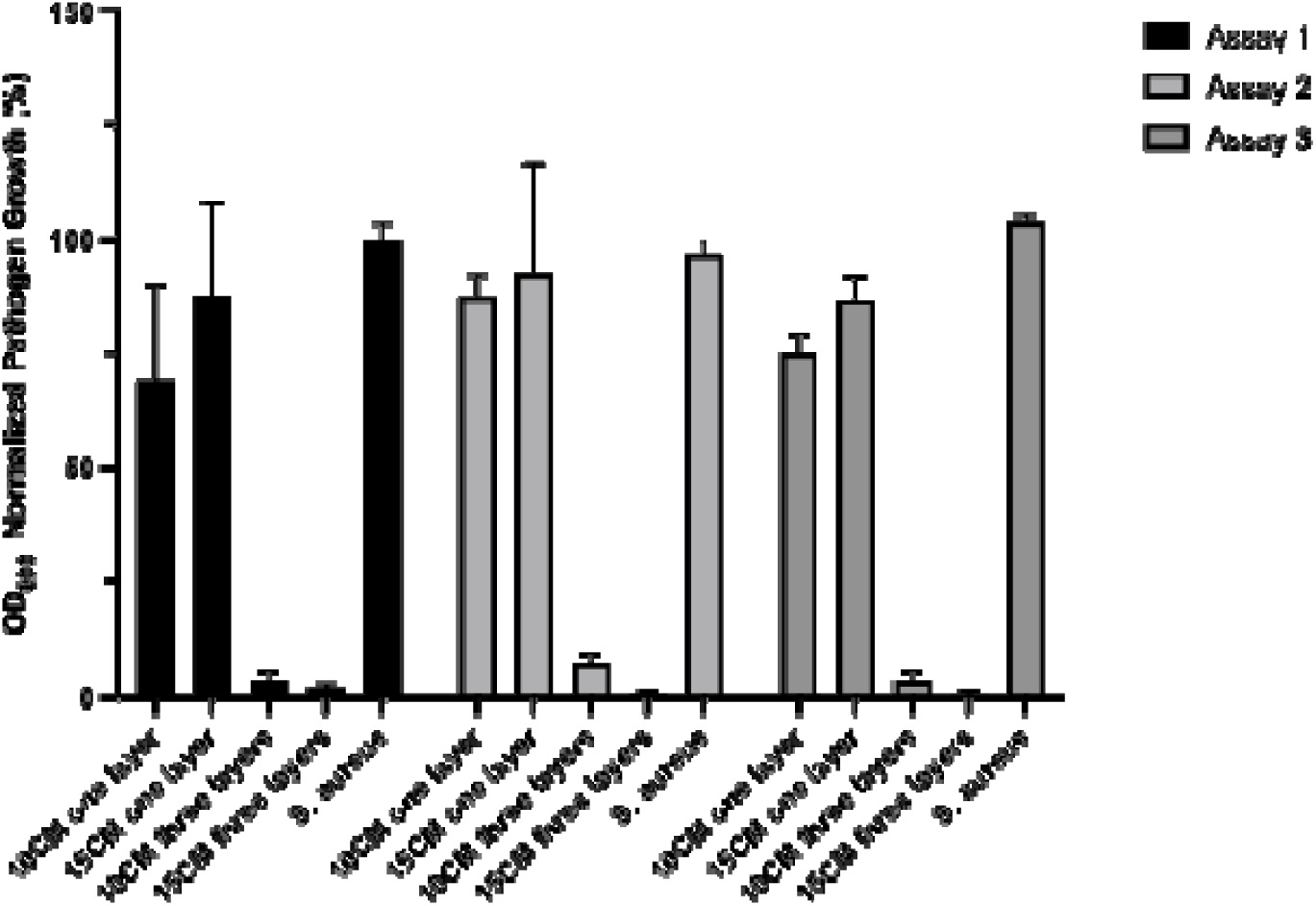
*In vitro* antimicrobial testing of a prototype spray device using εPLL (10 mg/mL) and HA (1 mg/mL) against *S. aureus*. Tested spray parameters included single vs. 3 coating layers and distances of 10 cm vs. 15 cm. Data are presented as means ± standard deviation from 3 biological replicates, and each biological replicated was measured with at least three technical replicates. Bacterial growth was normalized to *S. aureus* grown alone.

Results indicated that the number of spray layers significantly influenced the coating’s antimicrobial performance. Increasing the spray passes (spray from left to right) from one to three layers (3 passages) enhanced the antimicrobial efficacy against *S. aureus* ATCC 25923 (Figure 3), suggesting that multiple spray passages improve coating performance. With a total volume of 2.5 mL per polymer solution, three layers or even more could be applied. When assessing the influence of spray distance, it was observed that coatings produced at 10 or 15 cm exhibited strong antimicrobial activity against *S. aureus* (Figure 3).

After determining the optimal number of spray layers and spray distance, the polycationic εPLL was replaced with another polycation, namely Polyarginine 30 (Polymer chain of 30 arginine residues; PAR30), to comply with GMP standards. Indeed, PAR30 can be synthesized using solid-phase peptide synthesis (SPPS), a process that makes it easier to comply with GMP standards than the production of εPLL, which is produced from bacterial fermentation ^31,32^. Different concentrations of PAR30 and HA were evaluated *in vitro* against *S. aureus* ATCC 25923 and *Pseudomonas aeruginosa* ATCC 27853. Initially, PAR30 concentration was varied at 3, 5, and 10 mg/mL while HA was fixed at 3 mg/mL, leading to formulations referred to as 3P3H, 5P3H, and 10P3H. Tests were conducted at spray distances of 10 cm and 15 cm (Figure 4). Results showed that increasing the polycation concentration improved antimicrobial activity, particularly against *P. aeruginosa*. However, even the highest concentration (10P3H) did not exhibit strong activity against both bacterial strains.

**Figure 4.**
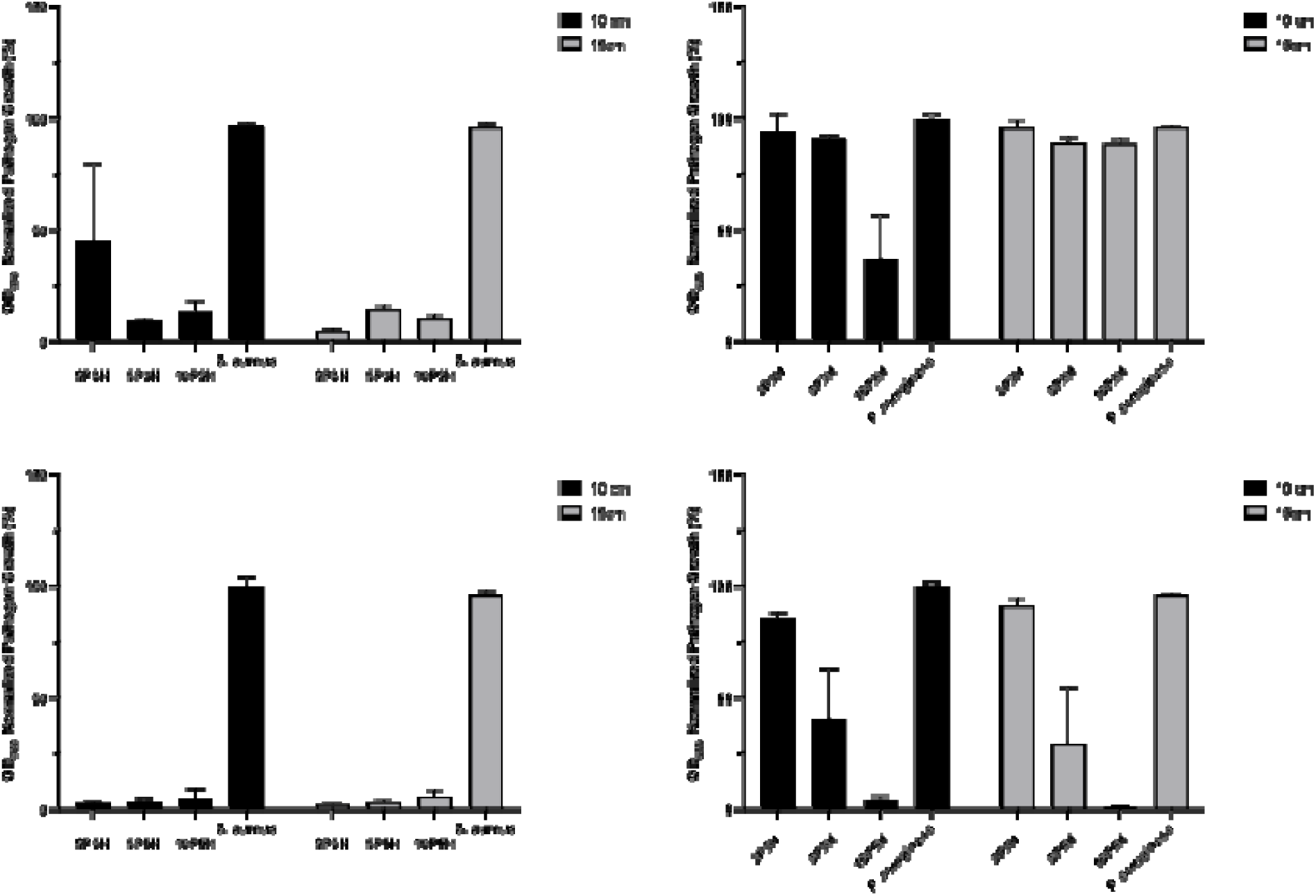
*In vitro* antimicrobial activity of PAR30 (3/5/10 mg/mL) with HA (3 or /5 mg/mL) against *S. aureus* and *P. aeruginosa* at 10 cm and 15 cm spray distances. Data are presented as means ± standard deviation from 3 biological replicates, and each biological replicate was measured with at least 3 technical replicates. Bacterial growth was normalized to *S. aureus* and *P. aeruginosa* grown alone.

To address this limitation, the HA concentration was increased from 3 mg/mL to 5 mg/mL while maintaining the same PAR30 concentrations, resulting in new formulations: 3P5H, 5P5H, and 10P5H. These were tested on optical glass slides at spray distances of 10 cm and 15 cm (Figure 4).

Increasing the polycation concentration with HA fixed at 5 mg/mL led to greater antimicrobial efficacy against both *S. aureus* and *P. aeruginosa* at each spray distance (Figure 4). Optical density measurements indicated that the 10P5H formulation yielded the most effective performance, supporting its selection as the optimal candidate for the final spray device (> 90% growth reduction). To assess whether the solutions could withstand sterilization, PAR30 and HA solutions were steam-sterilized and then tested in bacterial challenge assays as described above. Both solutions retained antimicrobial activity after autoclaving, resulting in a several-log reduction in bacterial load (Supplementary Figure S2). Overall, these findings support 10P5H as the lead formulation and suggest that the coating solutions can withstand autoclaving without loss of antimicrobial performance.

### In silico study of interactions in the PAR30/HA complex

To elucidate on the molecular basis of polycation/polyanion bilayers, molecular dynamics (MD) simulations were employed to investigate the intermolecular interactions between polycations, polyanions, and the solvent. The approach provides a deeper understanding of their behavior and stability in a simulated environment. We simulated the dynamics of the PAR30/HA complex in aqueous solution with a 150 mM concentration of sodium chloride. The interaction analysis revealed that PAR30 and HA interact primarily through polar and ionic interactions, whereas ionic interactions with sodium and chloride ions are less frequent for both HA and PAR30. The hydrophobic interactions between PAR30 and HA are also expected to be rare. The detailed analysis of position-specific interaction frequencies in PAR30 chains highlighted that certain residue, particularly the 1st residue of arginine, shows a high frequency of polar and ionic interactions (Supplementary Figure S3). In contrast, others, like the 11th and 23rd residues, consistently exhibit lower interaction frequencies across chains. These findings highlight the non-uniform contribution of individual residues to complex formation and underscore the importance of sequence position and structural context in governing PAR30–HA interactions.

### Supramolecular Coating Characterization

Building upon the observed antimicrobial efficacy of the 10P5H formulation, subsequent efforts focused on assessing its supramolecular structure and surface characteristics of the resulting coatings. Analyses were conducted on both glass slides and titanium (Ti40) to determine morphology, composition, and uniformity, linking biological performance to coated material properties.

Raman spectroscopy was applied to evaluate the chemical composition and structural features of the 10P5H coatings deposited on the Ti40 alloy and glass substrates, in comparison with their uncoated counterparts as well as with the reference PAR30 and HA (Figure 5). Spectra were collected over the 200–4000 cm^−1^ range. In the coated samples, weak low-frequency bands below approximately 400–600 cm^−1^ were observed for the Ti40 substrate and are attributed to the (Ti–O) and (Ti–N) stretching modes from native oxide or nitride layers. The glass substrate typically exhibits broad Si–O–Si network bands centered near 450–500 cm^−1^ and 1050–1100 cm^−1^; however, these substrate related signals were not clearly distinguishable due to the low Raman scattering efficiency of the inorganic phases and the dominance of the polymeric coating signals (Figure 5). Distinct diagnostic bands of the 10P5H coatings were clearly observed and are summarized in the accompanying Table 1. The band at ∼964 cm^−1^ is assigned to (C–O–C) stretching mode, indicative of ether linkages in HA PAR30 ^33^. The region around ∼1090 cm^−1^ corresponds primarily to (C–O) stretching of sugar/alcohol groups in HA, with possible contributions from C–C and C–N stretching characteristic of polyarginine. The band near ∼1330 cm^−1^ is attributed to (C–N) stretching and δ(CH /CH) bending, consistent with amino and alkyl groups present in polyarginine and HA ^34^. A further band around ∼1456 cm^−1^ corresponds to δ(CH /CH) bending vibrations, confirming the presence of polymer alkyl chains ^35^. The amide I band at ∼1684 cm^−1^, assigned to (C=O) stretching, is indicative of peptide bonds from polyarginine and/or carboxylate groups from HA, thus demonstrating the coexistence of both polymeric components ^36^. In the high wave number region, strong bands at ∼2930 cm^−1^ correspond to (C–H) stretching of CH /CH groups in the organic coating, while the region between ∼3200–3400 cm^−1^ is assigned to (O–H) and (N–H) stretching: specifically, ∼3200 cm^−1^ is attributed to strongly hydrogen bonded O–H groups in HA, ∼3350 cm^−1^ to predominantly N–H (amide/amine) with minor O–H contribution. Overall, the observed Raman spectral features confirm the successful deposition of the 10P5H coating and the presence of both HA and polyarginine components on both Ti40 and glass substrates. The absence of these diagnostic polymer-related bands in the uncoated Ti40 and glass substrates further confirms that the detected features arise specifically from the deposited biopolymer film.

**Figure 5.**
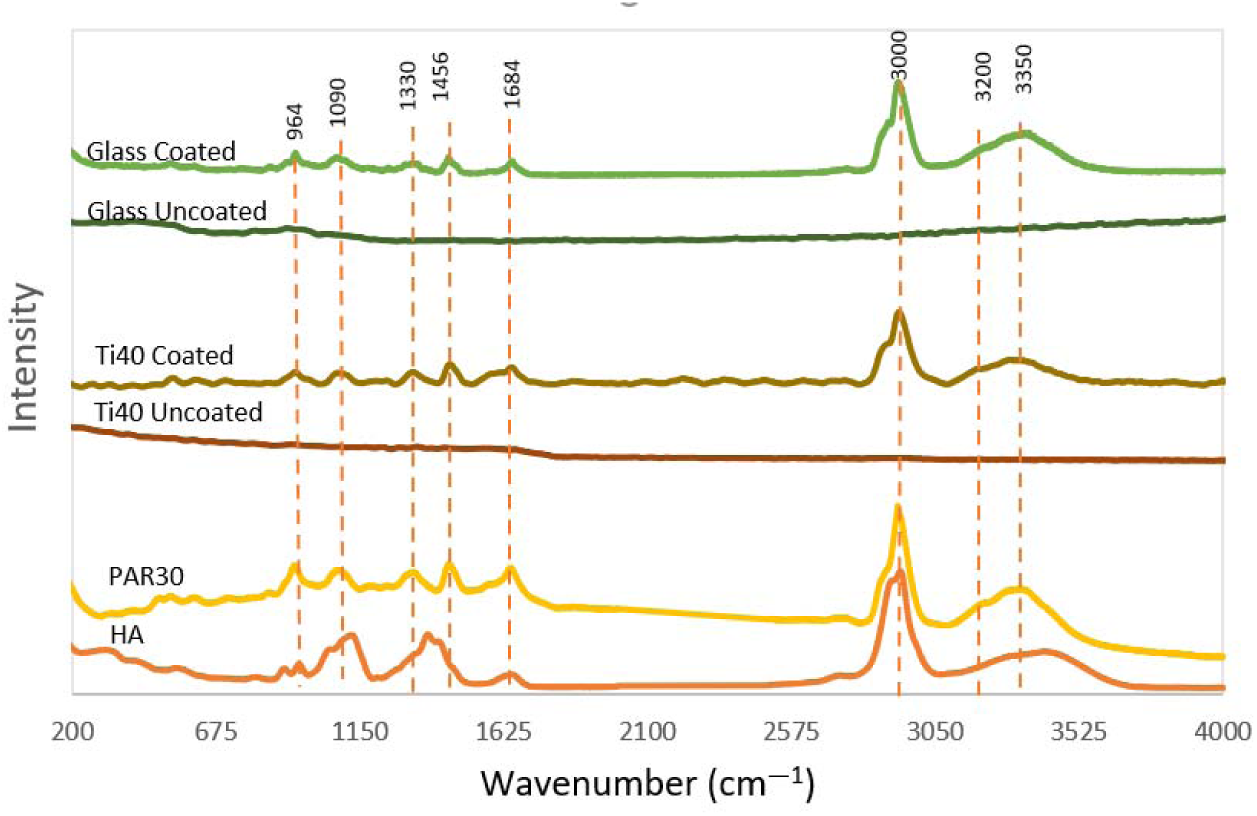
Raman spectra of uncoated and coated substrates (Glass and Ti40) compared with reference materials PAR30 and HA. Characteristic absorption bands are marked at 964, 1090, 1330, 1456, 1684, 2930, 3200, and 3350 cm□¹, corresponding to vibrational modes of C–O, C–N, C–H, C=O, and O–H functional groups

**Table 1:** Raman peak assignment of 10P5H coatings. Peak positions and tentative assignments according to references.

| Approx. position ($\text{cm}^{-1}$) | Probable functional group | Explanation |
| --- | --- | --- |
| ~964 | $\nu(\text{C–O–C})$ | Ether/ester bonds in (HA) |
| ~1090 | $\nu(\text{C–O})$ | Characteristic of sugar/alcohol groups in HA; possible polymer backbone vibrations |
| ~1330 | $\nu(\text{C–N}) / \delta(\text{CH}_2/\text{CH}_3)$ | Alkyl or amino groups in poly-arginine and HA |
| ~1456 | $\delta(\text{CH}_2/\text{CH}_3)$ | Methyl/methylene bending vibrations in polymers |
| ~1684 | $\nu(\text{C=O})$ | Peptide bonds in PAR30 or carboxylate groups in HA |
| ~2930 | $\nu(\text{C–H})$ | Stretching of $\text{CH}_2/\text{CH}_3$ groups in polymers |
| ~3200 | $\nu(\text{O–H, strongly H-bonded})$ | Hydroxyl groups in HA (strongly hydrogen-bonded) |
| ~3350 | $\nu(\text{N–H})$ (main) / $\nu(\text{O–H})$ (minor) | Amide or amine groups (PAR30), minor O–H |

Focused ion beam scanning electron microscopy coupled with energy dispersive spectroscopy (FIB–SEM/EDS) was performed to examine the morphology and elemental composition of the 10P5H coatings on glass and Ti40 substrates at magnification of 250× and compared with uncoated glass and Ti40. As shown in Supplementary Figure S4_I A, the glass-coated surface exhibited heterogeneous morphology with branched or dendritic structures, indicating a non-uniform deposition pattern with irregular grains and network-like growth. The Ti40-coated surface (Supplementary Figure S4_II A) displayed similar interconnected dendritic morphology with branched and irregular grains, highlighting the influence of substrate characteristics on coating nucleation and growth. Cross-sectional FIB images (Supplementary Figures S4_I B and S4_II B) revealed continuous coating layers with varying thicknesses on both substrates, confirming successful deposition. Quantitative analysis showed that the coating thickness was 3 ± 1 µm on the Ti40 substrate, and 3 ± 0.4 µm on the glass substrate.

The FIB cross-sections demonstrated good adhesion of the 10P5H coatings to both substrates. However, differences in layer uniformity and microstructural density between Ti40 and glass highlight the influence of substrate type on coating morphology and structural consistency.

**Energy-dispersive X-ray spectroscopy (EDS)** analysis (Supplementary Figure S5 and Supplementary Table S2) was used to assess the surface elemental composition of 10P5H-coated and uncoated glass and Ti40 substrates. The coated glass samples exhibited nitrogen (27.0 at.%), oxygen (25.5 at.%), sodium (19.0 at.%), and chlorine (28.5 at.%) as the predominant elements, consistent with the presence of the polymeric coating layer and residual NaCl from the buffer solution. In contrast, the uncoated glass substrate showed a homogeneous composition dominated by oxygen (61.0 at.%) and silicon (24.0 at.%), with minor amounts of sodium (7.0 at.%), magnesium (2.0 at.%), potassium (0.3 at.%), calcium (3.8 at.%), and tin (1.5 at.%). The larger variability observed in the coated glass samples reflects the heterogeneous nature of the deposited layer. For the Ti40 substrate, the coated samples contained nitrogen (27.0 at.%), oxygen (20.0 at.%), chlorine (10.5 at.%), and titanium (42.5 at.%), confirming the presence of the coating on the surface, while the detected titanium signal also originates from the underlying substrate. The pronounced differences between individual spectra indicate a non-uniform coating distribution across the Ti40 surface. By comparison, the uncoated Ti40 substrate consisted predominantly of titanium (89.0 at.%) with a minor oxygen contribution (11.0 at.%), attributable to the native oxide layer.

### Assessment of spray coatings using agar-based models

To better evaluate the coating under conditions representative of a wound environment, we sought an alternative to glass slides, which may not accurately simulate such conditions. Agar plates, with their moist and nutrient-rich surface, were chosen as a more physiologically relevant substrate that better mimics a wound bed compared to the previous surfaces tested above (e.g., glass and titanium). The 10P5H spray coating was therefore applied directly onto agar plates, followed by the deposition of standardized bacterial inoculum drops to assess antimicrobial efficacy (Figure 6). The coated agar plates were tested against *P. aeruginosa ATCC 27853*, *S. aureus ATCC 25923*, and *E. coli ATCC 25922* (Table 2). Significant log reductions in viable bacterial counts were observed (up to 7 log reduction). To ensure that the Tris–NaCl buffer (10–150 mM, pH 7.4) used to prepare the PAR30 and HA polymer solutions did not affect bacterial growth, buffer-only sprays were tested as additional controls and showed no inhibitory effects. Likewise, HA alone exhibited no antibacterial activity, confirming that the observed effects were solely attributable to the antimicrobial peptide. Notably, spraying PAR30 alone yielded similar log reductions, further demonstrating the peptide-driven antimicrobial mechanism.

**Figure 6.**
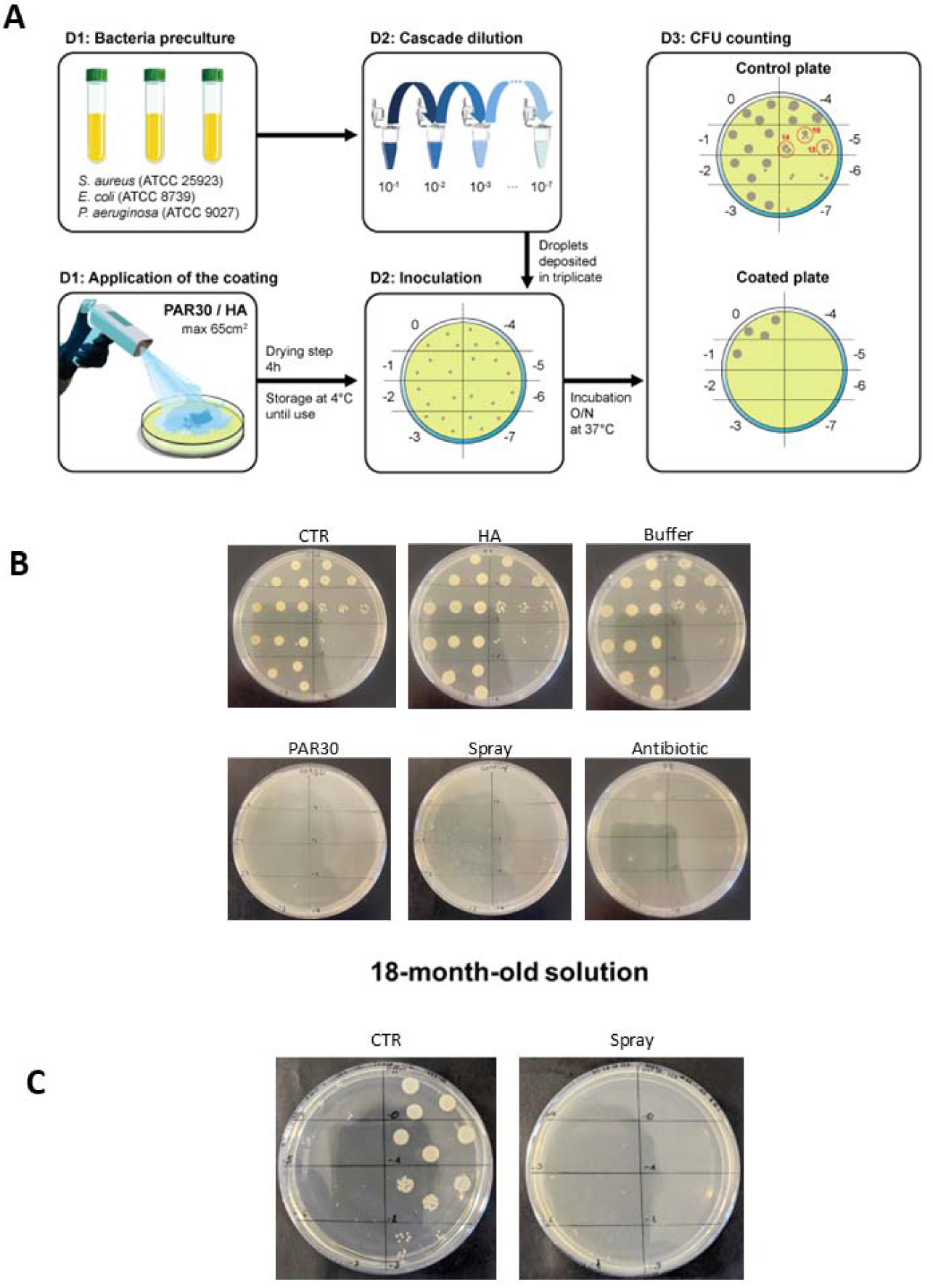
Antimicrobial evaluation of PAR30 peptide and HA coatings applied by spray on MH agar plate. (A) Schema drop plate method and bacterial enumeration (B) Representative agar plates showing *S. aureus* bacterial growth on (CTR) uncoated control, (Buffer) Tris-NaCl 20mM used to dissolve the polymers, (Spray) plate sprayed with polycation/HA, (PAR30) plate sprayed with AMP peptide only, and (HA) plate sprayed with hyaluronic acid only, Antibiotics (AB, Cefotaxime + Tetracycline) were also sprayed to have a comparison, illustrating the differential inhibition of microbial growth depending on the treatment. (C) Prefilled syringes were kept for 18-months in fridge and tested subsequently with exponential preculture of *S. aureus* to assess the efficacy of the coating after long-term storage.

**Table 2.** Summary of results from the drop plate method. The spray coating demonstrates antimicrobial activity against both Gram-positive and Gram-negative bacteria, effectively inhibiting growth at concentrations up to at least 10□ CFU/mL.

| Coating | Microorganisms | PAR30 |
| --- | --- | --- |
| PAR30 (10 mg/mL) /HA (5mg/mL) | *S. aureus* ATCC 25923 | 7 (+/- 1 log) |
|  | *E. coli* ATCC 25922 | 7 (+/- 1 log) |
|  | *P. aeruginosa* ATCC 27853 | 8 (+/- 1 log) |

Prefilled syringes with the polymer formulations were stored at 2–8 °C for 18 months to assess the effect of long-term storage on coating performance. After storage, the syringes were subjected to bacterial challenge assays identical to those used for freshly prepared samples. Similar log reductions in viable bacterial counts were observed for stored and freshly prepared syringes, indicating that prolonged storage did not adversely affect the antibacterial efficacy of the polymer coatings. These findings confirm that the coating is effective against both Gram-positive and Gram-negative bacteria. The approach offers notable advantages in simplicity and rapidity, enabling efficient antimicrobial performance under conditions that realistically approximate bacterial contamination.

### *In vitro* cytotoxicity

To assess the biocompatibility of the spray coating for biomedical applications, *in vitro* cytotoxicity testing (Alamar blue assay) was conducted on mouse embryonic fibroblast cell line Balb/3T3, in accordance with ISO 109935:2009, the international standard for evaluating the biological response of mammalian cells to medical device materials. Optical glass slides were spray-coated with the 10P5H formulation and evaluated using the direct contact method described in ISO 109935. According to ISO 109935 criteria, a reduction in cell viability of ≥ 30% signifies cytotoxicity. Cell viability assays confirmed that the 10P5H-coated glass slides supported cell viability above 70% after 24 h, like non-coated controls, and well above the cytotoxicity threshold. Also, no significant difference in fluorescence between coated and control groups was noted, thus validating the absence of cytotoxic effects (Figure 7). Overall, these results confirm that the spray-applied 10P5H coating is biocompatible *in vitro*, supporting its suitability for subsequent *in vivo* evaluation of antimicrobial efficacy and host inflammatory response.

**Figure 7.**
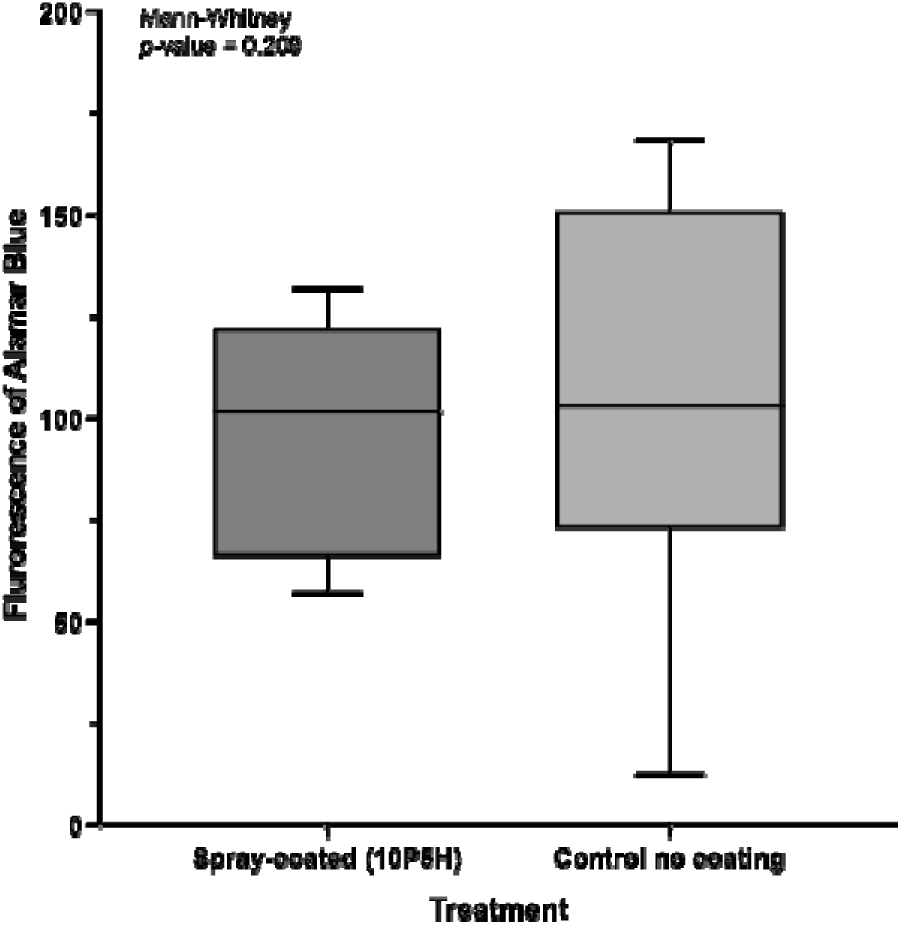
Spray toxicity on BALB3T3 during 24h incubation. Each experiment was performed in octuplicate and repeated 4 times. The statistical analysis used is the Mann-Whitney test, showing no statistically significant differences between groups (*p*= 0.209)

### *In vivo* mice model

#### Coating safety: Host inflammatory reactions and associated protein detection

Following the *in vitro* biocompatibility assessment on fibroblast cells, the 10P5H coating formulation was evaluated *in vivo* using SKH1 nude mice to assess potential inflammatory responses. Three groups were tested in this experiment, with two animals per group. Two full-thickness wounds (6 mm in diameter) were created on the dorsal skin of mice in the first two groups. The first group received the 10P5H spray treatment immediately after wounding, while the second group remained untreated. A third group, comprising non-wounded and untreated mice, served as the baseline control. On day 1, the coating was applied to the wounds of the treated group. After 3 h, the wounds in both the treated and untreated groups were stained using the IVISense MMP 750 FAST fluorescent probe, and all three groups were imaged via bioluminescence for immune monitoring of the host inflammatory response. Cytokine levels, specifically interleukin6 (IL6), were quantified at 24 h and 48 h post-treatment to further evaluate systemic and local inflammatory activity (Figure 8). Bioluminescence imaging revealed an initial increase in inflammatory activity at 3 h post-wounding in both the treated and untreated groups compared with the control, indicating a normal early inflammatory response to tissue injury. At 24 h, inflammation levels decreased markedly across all groups, and no significant differences were observed between the spray-treated and untreated wounded mice (Figure 8, B). IL6 quantification confirmed these observations: elevated cytokine concentrations were detected at 24 h in both wounded groups relative to controls, with no significant difference between the treated and untreated conditions. By 48 h, IL6 levels in both wounded groups declined sharply, approaching those of the control group (Figure 8, C). Collectively, these results indicate that the 10P5H coating does not exacerbate inflammatory responses and is well tolerated *in vivo*, supporting its biocompatibility for further preclinical evaluation.

**Figure 8.**
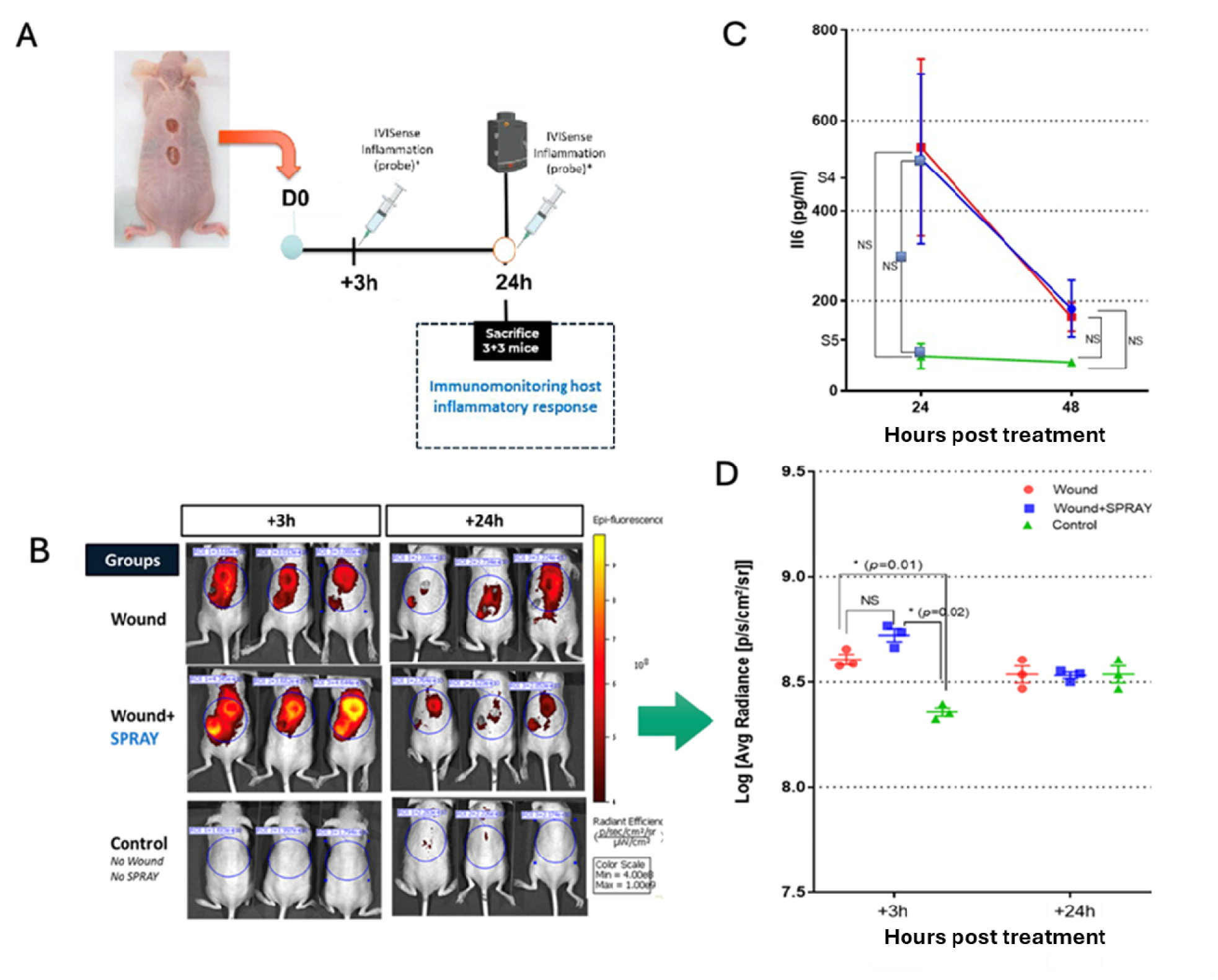
Immunomonitoring of host inflammatory response. **(A)** Schematic overview of the experimental protocol for assessing inflammatory host response following wounding and spray application administration. **(B)** Whole-body fluorescence imaging of mice using the IVIS Spectrum following intravenous administration of IVISense MMP 750 FAST, shown at 3 and 24 hours post-treatment (scale: p/s/cm²/sr). **(C)** Quantification of cytokine (IL-6) protein levels at 24 and 48 hours post-treatment. **(D)** Quantitative analysis of *in vivo* probe fluorescence signals (average radiance) comparing wound, wound and spray, and control mouse groups.

### In vivo antimicrobial activity

Bacterial proliferation and biofilm formation were assessed in a murine wound infection model using a bioluminescent strain of Methicillin-resistant *S. aureus* (MRSA: Xen31). Wounds were treated with 10P5H spray coating or left untreated as controls. After inoculation with bacteria and compress application, bacterial colonization was monitored by bioluminescence imaging and CFU counting (Figure 9 I).

**Figure 9.**
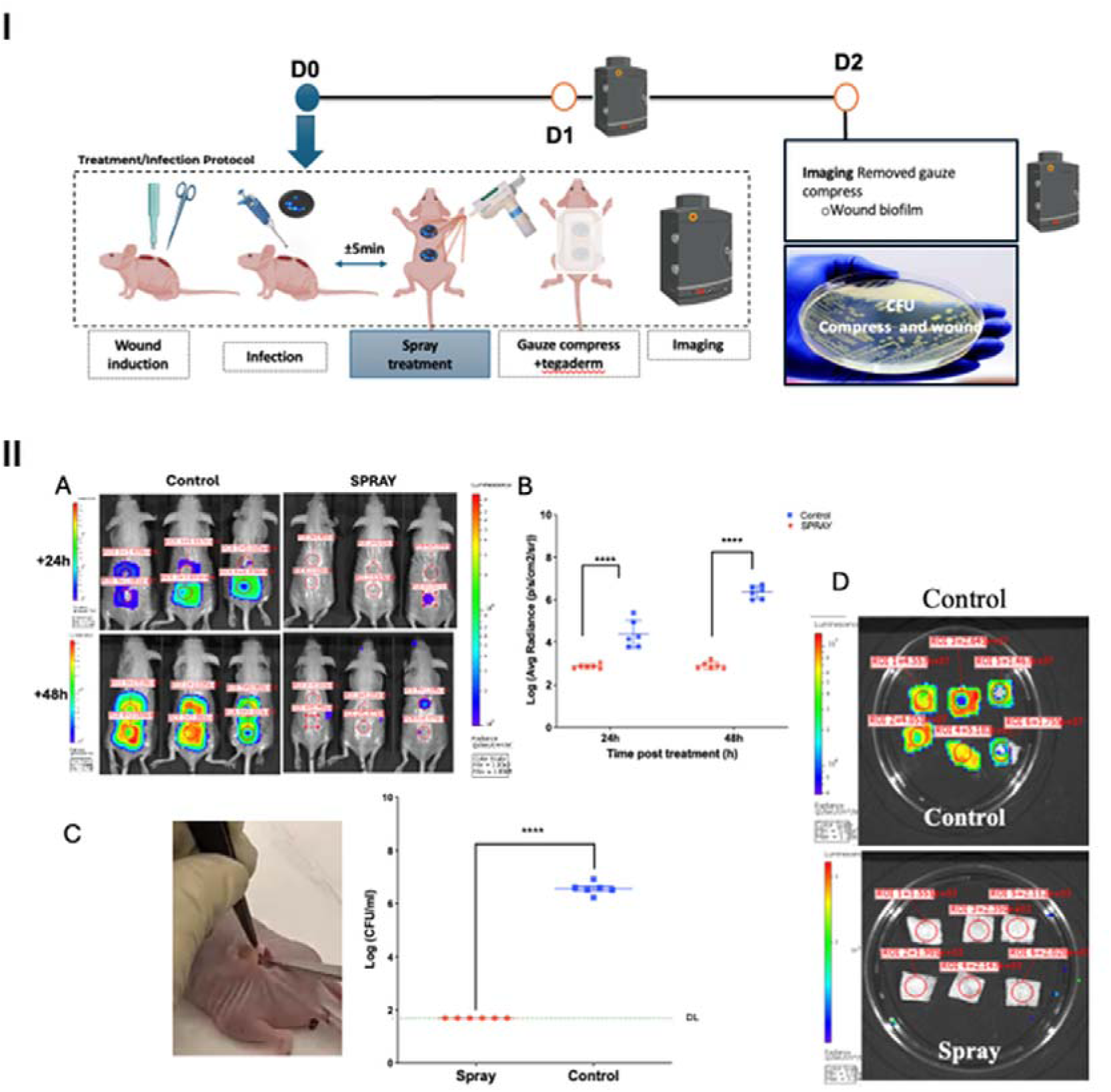
Evaluation of spray-coating efficacy in preventing bacterial contamination in a wound model. **(I)** This figure illustrates the experimental procedure used to assess the effectiveness of spray coating in preventing bacterial colonization in wound infection models. Bacterial suspensions were applied to either spray-coated or uncoated (control) wound sites to evaluate the coating’s ability to inhibit colonization. **(II)** monitoring of bioluminescent MRSA (xen31) in wound and gauze at 24h and 48h post-treatment spray/infection. (A) representative in vivo bioluminescent imaging of bacterial signals during implant infection. (B) quantification of in vivo bacterial bioluminescent signals presented as mean average radiance ([p/s/cm²/sr] ± standard error of the mean) on a logarithmic scale. Statistical analysis was performed using two-way ANOVA (\*\*\*\**p* < 0.0001). (C) quantification of bacterial load in compress gauze at 48h post-infection. Statistical analysis was performed using an unpaired test (\*\*\*\**p* < 0.0001). (D) bacterial bioluminescence and load in gauze at 48h post-treatment/infection. Representative bioluminescent imaging of bacteria in compress gauze after removal.

Bioluminescence imaging performed using the IVIS Spectrum system showed that bacterial signals remained very low in spray-coated wounds compared to uncoated controls during the first 48 hours post-infection (Figure 9 II, A and B). In untreated wounds, a strong bioluminescent signal was observed, indicating significant bacterial growth and biofilm formation. Quantification of bioluminescent intensity confirmed a significant reduction in bacterial load in the sprayed group compared to controls (two-way ANOVA, ****p < 0.0001). Bacterial burden at the wound site was further assessed by excising the tissue at 48 hours post-infection and performing CFU counts. Wound homogenates from the spray-treated group showed significantly fewer CFUs than those from the untreated group (Figure 9 II, C). Additionally, gauze compress covering the wound sites were analyzed for bacterial contamination. Bioluminescence imaging of the compresses revealed that dressing materials from the spray-treated wounds exhibited minimal or no detectable bacterial signals, whereas compresses from the untreated group were heavily contaminated. Quantification confirmed a statistically significant reduction in bacterial load on the gauzes from the coated group (Figure 9 II, D). Overall, the application of the antimicrobial spray coating effectively prevented bacterial colonization, reduced bacterial load at the wound site, and protected the dressing materials from contamination during the early stages of wound infection.

### Long-term effect on healing and wound contamination

To investigate the effect of spray treatment on bacterial colonization and wound repair, mice were assigned to two groups: control (wound only) and treated (wound plus 10P5H spray). Animals were sampled at multiple points for bacterial contamination, and wound closure was monitored throughout the study period (Figure 10, A). Bacterial burden around the wounds was quantified using non-selective UriSelect™ 4 agar and selective *Staphylococcus* Chromogenic Agar. Spray-treated wounds exhibited a significant reduction in total bacterial counts relative to controls at days 4, 6, 8, and 10 after injury (Figure 10, B). Enumeration on Staphylococcus Chromogenic Agar further confirmed a lower staphylococcal load in the treated group (Figure 10, C). Analysis of wound closure revealed that spray application did not alter healing rates compared to the control group (Figure 10, D). Collectively, these data demonstrate that repeated spray treatment effectively reduces both the overall and staphylococcal burden in wounds, without affecting the healing process in a murine model.

**Figure 10.**
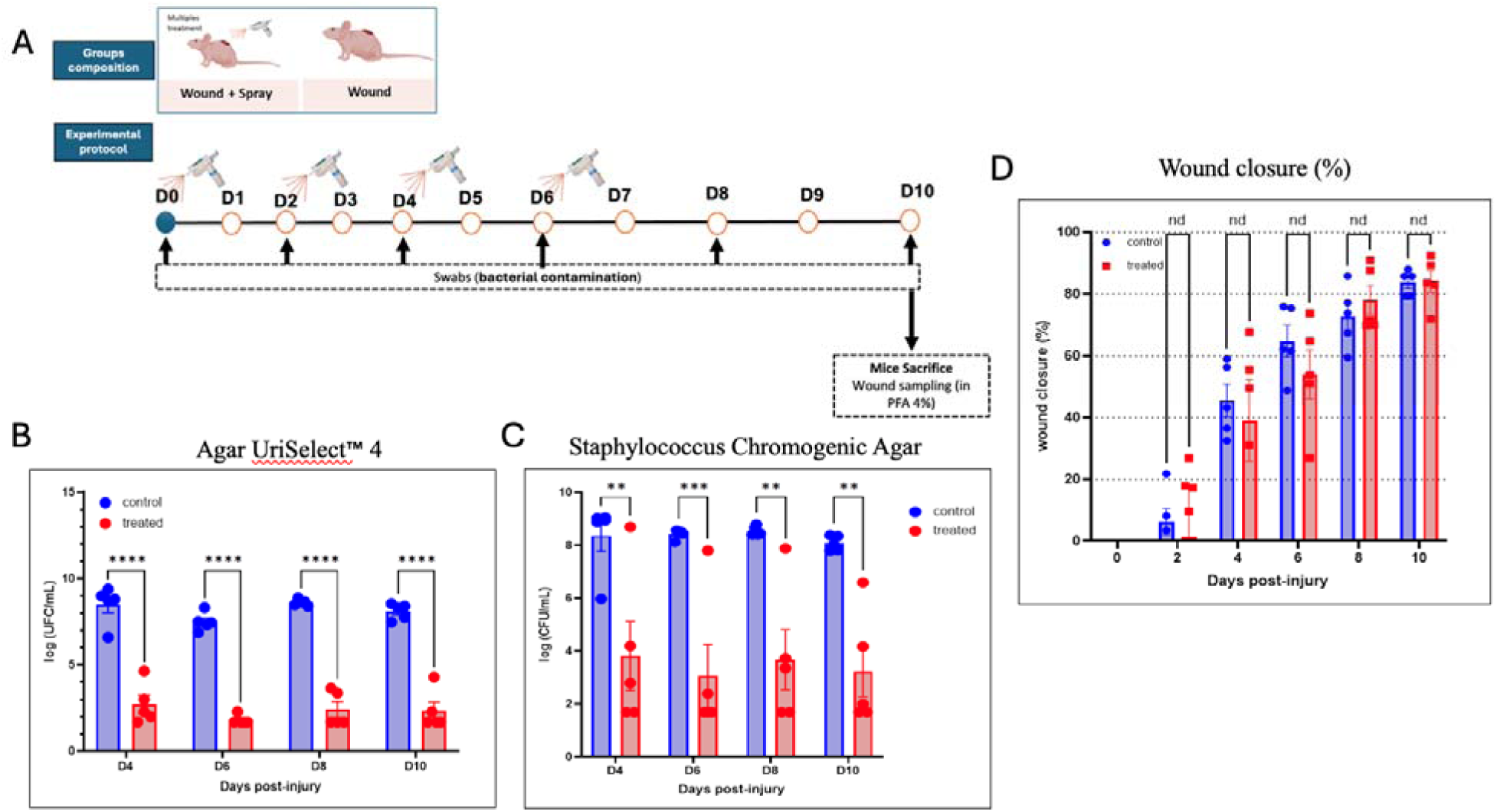
Evaluation of spray treatment effect on wound bacterial contamination and healing in mice. **(A)** Experimental protocol depicting control and treated (wound + spray) groups, swabbing schedule for bacterial analysis, and timeline for sacrifice and wound sampling. **(B)** Quantification of total bacterial load from wound swabs plated on Agar UriSelect™ 4 at days 4, 6, 8, and 10 post-injury. **(C)** Quantification of *Staphylococcus* spp. from wound swabs plated on *Staphylococcus* Chromogenic Agar at the same time points. **(D)** Percentage wound closure measured over 10 days in control and treated groups. Statistical analysis: Mann-Whitney test, \*\**p* < 0.01, \*\*\**p* < 0.001, \*\*\*\**p* < 0.0001.

### Behavioral effects of spray treatment on post-surgical hypersensitivity

To investigate the effect of 10P5H spray treatment on post-surgical pain and recovery, mice were divided into four groups: naive + saline, naive + HA/PAR, incision + saline, and incision + HA/PAR. The spray was applied twice during surgery, once 2 h after, and once 24 h after surgery, for a total of four applications. Behavioral and nociceptive assessments were performed throughout the study period to evaluate immediate reactions to spraying, as well as the development of mechanical, thermal, and cold hypersensitivity (Figure 11, A). We first assessed the immediate behavioral response to spray application. At 2 h post-surgery (after the third spray application) and at 24 h post-surgery (after the fourth application), incised animals exhibited licking and shaking behaviors in response to spraying. However, these responses did not differ between the “incision + HA/PAR” and “incision + saline” groups, indicating that HA/PAR did not produce greater irritant effects than saline (Figure 11, B) (F_3_,_92_ = 6.256, p = 0.0007; post-hoc: [incision + HA/PAR = incision + saline] > [naive + HA/PAR = naive + saline] at p<0.0001 after the third and the fourth application). We then evaluated the effects of HA/PAR treatment on nociceptive hypersensitivity. Saline-treated incision mice displayed mechanical hypersensitivity at days 2 and 4 post-surgery (von Frey, F□□,□□□ = 4.76, p < 0.0001), whereas “incision + HA/PAR” mice did not develop mechanical hypersensitivity (Figure 11, C). Similarly, “incision + saline” mice exhibited heat hypersensitivity at days 3 and 5 (Hargreaves, F□,□□= 30.68, p < 0.0001), with withdrawal latencies significantly shorter than those of HA/PAR-treated and non-incised controls (p < 0.001) (Figure 11, D). HA/PAR treatment completely prevented the development of heat hypersensitivity. Finally, cold sensitivity assessed with the dry ice test revealed significant hypersensitivity in “incision + saline” mice at days 1, 4, and 6 (F□, □□□= 17.84, p < 0.0001). Although HA/PAR-treated incision mice showed mild cold hypersensitivity, it remained significantly attenuated relative to saline-treated incision mice (p < 0.01) (Figure 11, E).

**Figure 11.**
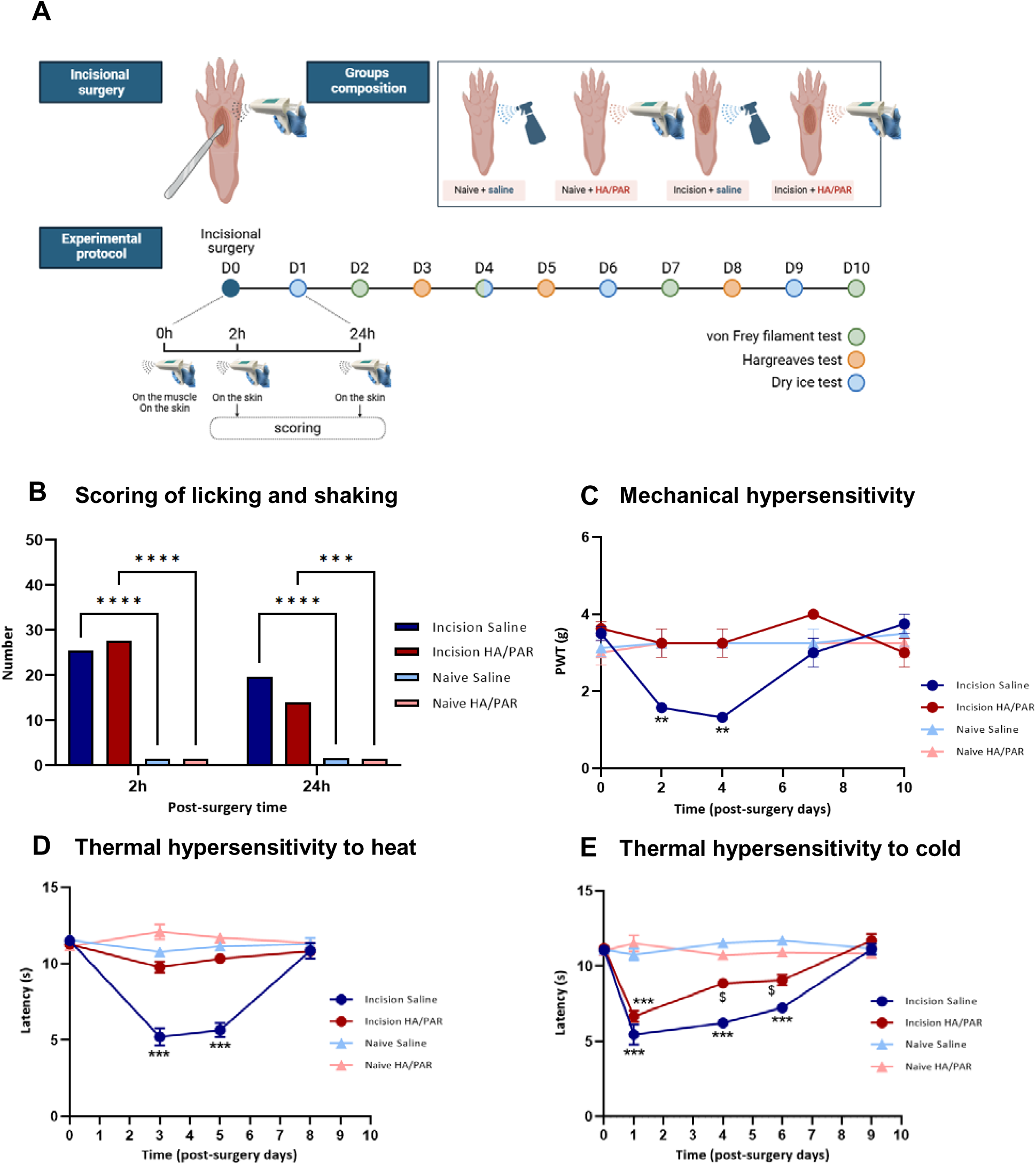
Behavioral effects of 10P5H spray treatment on post-surgical nociception. **(A)** Schematic representation of the experimental design showing the incisional surgery, naive and incision groups treated with either saline or 10P5H spray, and the timeline of behavioral testing until sacrifice. **(B)** Immediate behavioral responses to spray application assessed 2 h and 24 h post-surgery. Both incised groups displayed increased licking and shaking behaviors relative to non-incised animals; however, 10P5H treatment did not elicit greater nociceptive responses than saline, indicating good tolerability and absence of irritant effects. **(C)** Mechanical hypersensitivity evaluated using von Frey filaments. Saline-treated incision mice developed significant mechanical hypersensitivity at days 2 and 4, whereas 10P5H-treated incision mice did not. **(D)** Heat hypersensitivity assessed using the Hargreaves test. Saline-treated incision mice showed reduced paw withdrawal latencies at days 3 and 5 compared with HA/PAR-treated and non-incised controls. **(E)** Cold hypersensitivity assessed using the dry ice test. Saline-treated incision mice exhibited marked cold hypersensitivity at days 1, 4, and 6, while 10P5H - treated incision mice showed only mild and significantly attenuated responses. Data are presented as mean ± SEM. **\*\****p* < 0.01 and **\*\*\****p < 0.001 vs* Naive + Saline group; ^$^*p* < 0.01 *vs* Incision + Saline group.

Together, these results indicate that 10P5H spray triggers transient behavioral responses upon application but effectively prevents the development of mechanical and heat hypersensitivity and markedly attenuates cold hypersensitivity following paw incision, thereby improving recovery from post-surgical nociception.

### Effect of spray treatment on motor coordination in the beam-walking task

Motor coordination was assessed on the narrow 1.2 cm beam at day 7 post-surgery. No significant differences in traversal time were observed between the incised and non-incised groups (Figure 12, A). In contrast, incision significantly increased the number of slips, with saline-treated incision mice exhibiting an average of 8 slips across the three 100 cm trials compared with 2.3 slips in non-incised controls (F, = 9.356, p = 0.006). Application of the 10P5H spray tended to reduce the number of slips, although the overall effect did not reach significance (Figure 12, B). Analysis of the first 100 cm trial revealed a group × treatment interaction (F, = 4.708, p = 0.0416), with saline-treated incision animals displaying significantly more slips than all other groups (post-hoc p < 0.05). Notably, 10P5H-treated incision animals performed at levels indistinguishable from non-incised controls (Figure 12, C).

**Figure 12.**
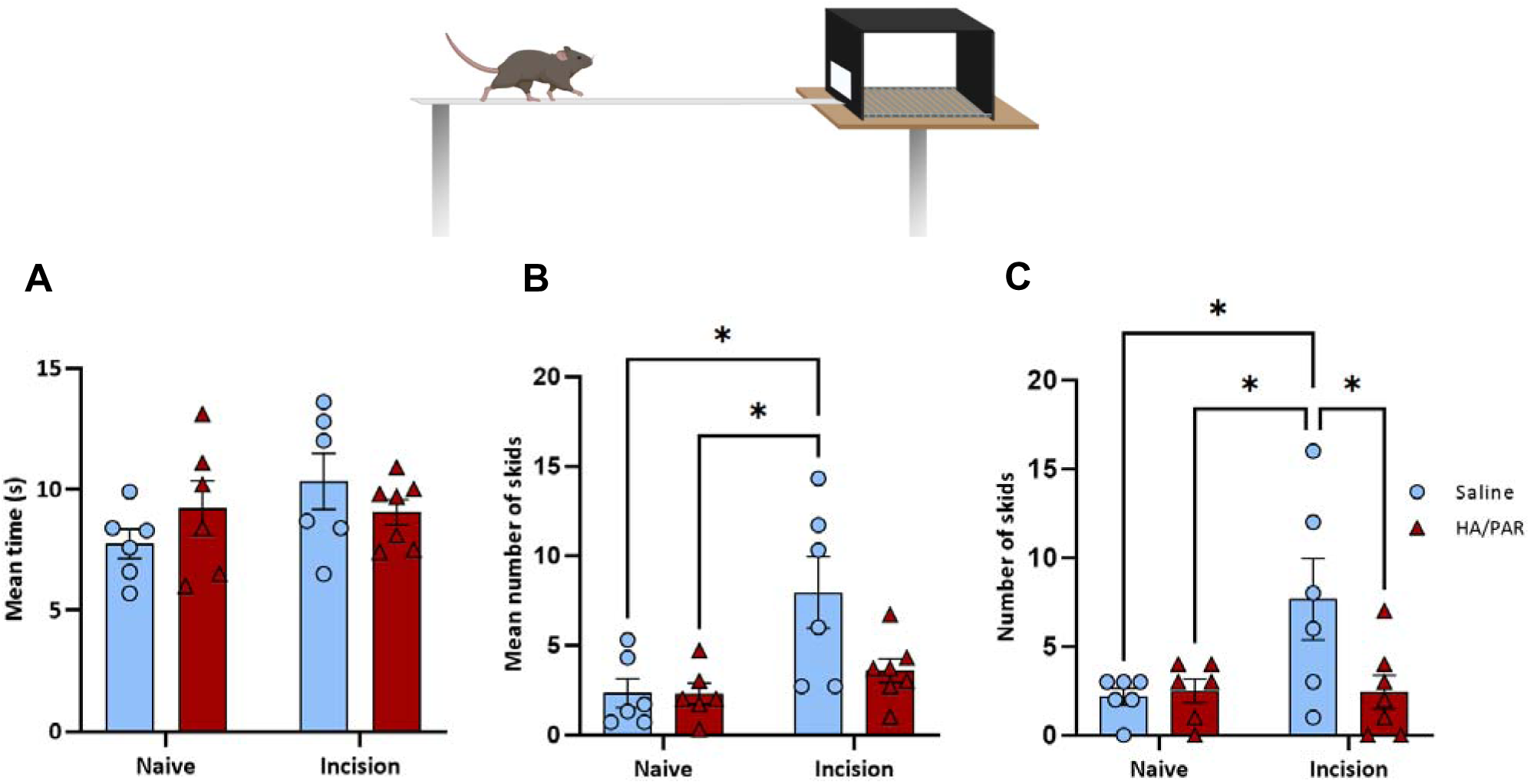
Effect of HA/PAR spray treatment on motor coordination in the beam-walking test. **(A)** Traversal time across the beam. No significant difference in traversal time was observed between incised and non-incised groups, irrespective of treatment. **(B)** Total number of slips across the three trials. Incised mice exhibited more slips than non-incised controls, with saline-treated incision mice showing the greatest impairment. **(C)** Mean number of slips during the first trial. Saline-treated incision mice displayed significantly more slips than all other groups, whereas HA/PAR-treated incision mice performed comparably to controls, indicating preserved motor coordination. Data are presented as mean ± SEM. \**p < 0.05 vs Naive + Saline group*.

These results indicate that while paw incision impairs fine motor coordination, leading to increased slipping on the narrow beam, 10P5H spray mitigates this deficit and restores motor performance to control levels.

### Safe and Sustainable by Design Assessment

We performed a simplified Safe-and-Sustainable-by-Design (SSbD) assessment to identify translational safety and sustainability hotspots associated with the biopolymer solution. The assessment covered hazard classification, occupational health and safety, human and environmental risk during the use phase, and a life-cycle assessment of the biopolymer solution. For the hazard classification, no cytotoxicity (Figure 7) and no host inflammatory reaction (Figure 8) were observed in vitro and in vivo studies, respectively. HA and PAR30 dissolved in water were tested separately using the lower tier OECD #442D test ^37^ keratinoSens-based approach. HA was found to be neither a sensitizer nor an irritant, whereas PAR30 showed an indication of potential irritancy (Supplementary Table S3), however this effect was observed only when PAR30 was tested alone and was not observed during the higher tier *in vivo* tests. Considering this effect was observed only for PAR30 tested in isolation and was not detected for PAR/HA combinations or in the higher-tier *in vivo* safety studies, suggesting that association with HA may mitigate the local irritancy of PAR30. Occupational risks during processing, assessed using StoffenManager, were low, with the lowest risk category assigned for both inhalation and skin contact/dermal uptake (Supplementary Table S4). Eye exposure during formulation was rate as “average”, supporting the use of appropriate eye protection, which is already required under ISO 7 cleanroom practice. For human health, a conservative inhalation-based use-phase scenario did not indicate a risk for either PAR or HA. Assuming complete aerosolization in a 35 m³ hospital room without ventilation and once-daily application for 5 consecutive days, the estimated exposure yielded a risk characterization ratio (RCR) of 0.05 for PAR, based on a surrogate derived no-effect level (DNEL), and a RCR of 0.89 for HA, based on a literature-derived no-adverse-effect reference value. Both values were below 1, indicating no expected human-health risk under this worst-case scenario. An environmental exposure screening based on worst-case release assumptions likewise did not indicate a concern (Supplementary Table S5). A simplified life-cycle assessment (Supplementary Table S6) nevertheless identified potential environmental hotspots associated with solution production, although these findings should be interpreted cautiously because the assessment relied on proxy datasets (Supplementary Table S7) and restricted system boundaries. Additional considerations did not reveal any points of concern (Supplementary Table S8).

## Discussion

In this study, we developed a spray-delivered hyaluronic acid/polyarginine (HA/PAR30) multilayer coating intended for *in situ* application on wounds and implantable or indwelling device surfaces. Using Raman spectroscopy and FIB-SEM, we verified the formation of continuous multilayer films on clinically relevant substrates (glass and titanium), and showed antibacterial efficacy was retained despite some thickness variability. Importantly, the coating reduced bacterial burden and prevented biofilm establishment while remaining cytocompatible and well tolerated *in vivo*. Notably, treatment also reduced post-surgical hypersensitivity without impairing motor coordination, supporting its potential as a practical, in situ, infection-prevention strategy for irregular wound geometries and device-associated settings. The significance of an *in situ*, conformal antibacterial coating becomes clear when considering the central role of biofilms in chronic wounds, particularly around implanted or indwelling medical devices ^38^.

Biofilms are structured microbial communities encased in extracellular polymeric substances (EPS) and show increased tolerance to host immune clearance and conventional antimicrobial therapies ^8^ . Biofilm-derived secreted products can directly impair cellular wound healing events, including fibroblast and keratinocyte proliferation and migration, while also promoting apoptosis in relevant cell types ^39^. In chronic wounds, biofilm aggregates formed by organisms such as *S. aureus* and *P. aeruginosa* ^5–7,40,41^ are associated with a relatively low-grade inflammatory response that impedes epithelial migration and granulation tissue ingrowth, thereby delaying wound closure ^39^.

This persistent inflammation, coupled with the physical barrier created by the EPS matrix, creates a microenvironment that favors chronicity and recurrent infection ^42–45^ and is associated with persistent activation of innate immune pathways such as TLR2 and NF-κB ^46^ . Irregular wound geometries create cavities that are hard to irrigate debride, and fill uniformly, which reduces therapeutic effectiveness and favors bacterial persistence and biofilm establishment. The development of antimicrobial coatings represents a promising strategy to prevent bacterial colonization, disrupt biofilm formation, and restore the wound healing process ^22,47^. By incorporating agents that targets bacterial membranes and/or disrupt the EPS matrix, antimicrobial coatings may prevent biofilm establishment ^48–50^.

Silver or similar metal ions are widely used as antimicrobial and antibiofilm agents ^51^ . Chegini et al., recently developed a hydrogel dressing combining nitric oxide (NO), silver nanoparticles, and the antibiotic ciprofloxacin (CIP) that demonstrated potent inhibition of bacterial growth and mature biofilms, while accelerating wound closure and improving tissue regeneration ^52^ . The efficacy of a silver oxynitrate-containing dressing against bacterial biofilms and the impact on wound healing have also been demonstrated ^53^. Multiple oxidation states of silver ions (Ag^1+^, Ag^2+^, and Ag^3+^), when incorporated into wound dressings, exhibit significantly enhanced antimicrobial activity against mature biofilms of *S. aureus* and *P. aeruginosa in vitro* and in murine infected wound models ^53^.

A sprayable collagen-chitosan hydrogel containing silver nanoparticles reduced *in vivo* bacterial load while accelerating healing and tissue regeneration ^54^. Similarly, a sprayable film made with nanocellulose and L-serine-modified silver nanoparticles demonstrated protection against bacterial infections, as well as improved healing and reduced inflammation^55^. While effective, accumulating evidence suggests that metal ions carry some risks, including potential cytotoxicity effects and the possibility of developing resistance or reduced effectiveness over prolonged use ^56^. Similar to antibiotics, the excessive use and misuse of silver compounds have increased the prevalence of silver-resistant microorganisms in the environment, posing a serious risk of widespread transmission, including through *sil* and *cus* operons-mediated resistance, and raising broader human health concerns ^57,58^.

In response to these risks, antimicrobial peptides (AMPs) have emerged as promising alternatives, particularly for topical wounds, delivered locally as degradable hydrogels or films, reducing systemic exposure ^59^. These AMPs, offer broad-spectrum activity against biofilms, a lower propensity for resistance, and additional benefits in promoting wound healing ^60,61^. Electrostatic interactions between cationic peptides and bacterial membranes have been shown to effectively neutralize pathogens ^24,62,63^, while materials such as hyaluronic acid promote tissue regeneration and reduce inflammation ^64^. HA layers on substrates further inhibit protein adsorption and resist bacterial adhesion due to their antifouling properties ^65^. Importantly, HA/Poly-epsilon-L-Lysine-based coatings have been shown to reduce infection risk on titanium and ultra-high-molecular-weight polyethylene while maintaining osteogenic activity ^66^.

On wounds, the synergy between antimicrobial agents and biocompatible materials can be used to maintain hydration and deliver sustained antimicrobial activity, which is essential for preventing device-related infections in clinical settings ^64^. In this context, the homopolycation polyarginine (PAR30) offers contact-killing activity through electrostatic interactions with bacterial membranes and exerts lower selective pressure for classical antibiotic resistance ^15,16^. Indeed, in a 30-day serial-exposure assay, PAR30 showed no detectable resistance development, consistent with a low propensity for resistance selection^16^ .

The HA/PAR30 multilayer coating technology has previously been applied to hernia meshes ^15^ and wound dressings^19^ through the LbL dipping method, demonstrating strong antibacterial activity against wound pathogens in both *in vitro* and *in vivo* studies, preventing bacterial colonization and promoting healing. In this study, HA/PAR30 multilayers were delivered via spray, and both Raman spectroscopy and FIB-SEM analyses confirmed the formation of continuous multilayer coating layers on glass and titanium substrates. Although the coating exhibited variable thicknesses, its deposition still maintained significant and potent antibacterial activity.

Beyond demonstrating efficacy, the translational potential of this platform also depends on its safety and sustainability profile during use and along its life cycle. From the SSbD perspective, the choice of biopolymers for antimicrobial and coating-forming activity ensures a local biodegradation to the amino acids and disaccharides, thereby minimizing potential release into the environment and downstream effect on flora and fauna ^67^. Moreover, this local degradability may help prevent sub-minimum inhibitory concentration accumulation of the antimicrobial components, as can occur with many biocides, and thereby reduce the risk of resistance development in hospitals and in the environment in general ^68^. The simplified SSbD assessment provides a rapid, early-stage overview and helped identify potential hotspots. Overall, the results indicate no red flags in the assessed safety evaluations; the potential irritancy of PAR30 alone should be monitored if the formulation is modified in the future. For the sustainability assessment, the main impacts of the solution were associated with glucose or ethanol production, which were used as proxies for PAR and HA. Sukumaran and Gopi (2021) and Grant (2012) reported that biologically derived polymer production has impacts originating from land use, which is consistent with our results and may be linked to the inclusion of the background system (raw material production, maize for glucose, and sugar beet for ethanol in the selected proxies) ^68,69^. The main challenge in the life cycle assessment (LCA) was the absence of inventory data for biopolymer production, requiring the use of proxies that may introduce uncertainty, as impacts can be highly dependent on factors such as technology, landfill performance ^68^ or organic material sources ^69^. However, a main goal of an early-phase SSbD assessment is to identify hotspots for further analysis and not provide final answers. Improved SSbD assessments in the next phase should focus on strengthening the analyses as more data become available, conducting a more precise and detailed inventory assessment for the LCA (e.g. modelling the ingredients with real data or choosing different proxies with adjustments), expanding hazard classification to include additional classes, and incorporating a socio-economic assessment. In addition, material efficiency has not been considered as the biopolymer solution is still in the innovation phase; however, optimizing the volume for the application is crucial to prevent unnecessary material use. Taken together, these early SSbD considerations support the broader translational promise of the coating while identifying priorities for further optimization.

The antimicrobial spray coating developed in this study addresses a critical unmet need in the management of chronic wounds and device-associated infection. The ability of the HA/PAR coating to form an ultra-thin, conformal film on a wide variety of substrates, including living tissues and implantable materials, offers a versatile platform for infection prevention across diverse clinical scenarios. Our findings demonstrate that this coating not only reduces bacterial loads significantly but also prevents biofilm establishment, a known driver of chronic infection and delayed healing. Physicochemical analyses, including Raman spectroscopy and FIB-SEM, further confirmed the intimate spatial association of the polycationic and polyanionic components, rather than their segregation into separate clustered islands. This supports the proposed electrostatic zip-like macromolecular structure of the coating.

A key advance of this association of biocompatible components (HA/PAR) is their synergistic combination of antimicrobial efficacy with anti-inflammatory and regenerative properties, supporting tissue repair while minimizing cytotoxicity. This was confirmed through comprehensive *in vitro* cytotoxicity studies and *in vivo* inflammatory marker monitoring, which showed no adverse host responses. This favorable safety profile holds promise for clinical translation, where biocompatibility is paramount. Importantly, our findings also demonstrated that HA/PAR spray coating prevents the development of mechanical and heat hypersensitivity, as well as cold hypersensitivity. HA is a therapeutic biomaterial that can reduce inflammation and pain ^23,70^. Recently, HA has been shown to directly inhibit the TRPV1 channel activity, decreasing nociceptor excitability and reducing behavioral response to noxious heat *in vivo* ^71^ . TRPV1 can also be antagonized by short arginine-rich peptides; indeed, Himmel et al., demonstrated that the synthetic hexapeptide R4W2 (RRRRWW-NH_2_) was a potent, stereoselective TRPV1 antagonist in adult rat dorsal root ganglion neurons, inhibiting capsaicin and resiniferatoxin-evoked Ca 2+ responses ^72^.

The LbL dip coating method is suitable for applying uniform and conformal films to an implant, but it is typically performed as a pre-operative step before the implant is brought into the operating room. In contrast, a portable applicator enables *in situ* coating of both wounds and implantable medical devices. From manufacturing and infection-control perspectives, conventional LbL dip baths constitute static liquid reservoirs prone to composition drift and environmental contamination, whereas the dual-cartridge spray format keeps the precursor solutions physically separated, facilitates sterile handling, and is more easily aligned with GMP requirements.

Additionally, the proposed spray system, is designed to be user-friendly, making it suitable for deployment across a range of healthcare settings, including resource-limited environments. Its ergonomic dual-nozzle mechanism enables simultaneous delivery of the coating components without the need for intermediate rinsing, streamlining administration and enhancing user compliance. Such ease of use is particularly important for the management of complex wounds or multi-material implants. The observed reduction in nociceptive hypersensitivity, alongside robust antimicrobial activity, suggests a dual benefit: improved infection control and potentially improved patient comfort and quality of life, which remains an under-emphasized aspect of wound management ^73^. Wound infection is well recognized to exacerbate pain, and pain is closely linked with psychological stress, and stress can, which in turn, impair the wound-healing process ^73^. Our findings suggest broader therapeutic potential beyond infection prevention, contributing to improved patient outcomes and faster recovery. Despite these promising outcomes for this prototype, large-scale clinical trials will remain necessary to validate its effectiveness and safety in diverse patient populations, particularly those with comorbidities common in chronic wound patients or in the case of invasive surgical procedures.

In summary, this innovative biopolymer spray coating represents a significant step forward in antimicrobial strategies for wound and device infection management, aligning with antimicrobial stewardship principles by reducing reliance on systemic antibiotics. Its adaptability, safety, and efficacy position it as a transformative technology with the potential to improve clinical outcomes, reduce healthcare costs, and mitigate antibiotic resistance development.

## Methods

### Polyelectrolytes and buffer

*Polycations:* poly (L-arginine hydrochloride) (PAR) from Alamanda Polymers, USA. PAR30 consisted of 30 arginine residues with a molecular weight (Mw) of 6.4 kDa. ε-poly-L-lysine (εPLL) was purchased from BIOSYNTH, USA, average molecular weight provided by the manufacturer was between 2000 to 4700 Da. *Polyanion:* Hyaluronic acid (HA, Mw = 144 kDa), was purchased from Lifecore Biomed, USA. *Buffers*: Buffers used to dissolve polymers were made by Tris base (Fisher bioreagents, USA) and Sodium chloride NaCl (Fisher bioreagents, USA). The buffer made for PAR and HA solution was 10 mM Tris base and 150 mM NaCl with distilled water. The buffer made for εPLL and HA solution was 20 mM Tris base and 20 mM NaCl with distilled water. Both buffers’ pH was adjusted to 7.4 by slowly adding 1 mol/L hydrochloric acid (VWR chemicals, USA). The solutions can be sterilized in an autoclave (121.4°C with a 20-minute plateau) without degradation or loss of antimicrobial properties.

### Substrates

*Glass slides.* 12 mm diameter optical cover glasses (Epredia, USA) were cleaned via a multi-step protocol: immersion in 2% Hellmanex (Hellma GmbH, Germany) and sonication for 30 min; rinsing with distilled water followed by immersion in 1 M HCl for 15 min; final sonication in 70% ethanol (VWR, USA) for 30 min. Slides were stored in 70% ethanol at room temperature. *Medical Grade Titanium.* Titanium disks (13 mm diameter, 6 mm thickness; ACNIS, France) were used to assess the applicability of the coating system to metallic implants. Disks (size) were cleaned by 10 min sonication in 70% ethanol and stored in sterile containers.

### Spray coating protocol

The polymer solutions were added to the left side (polyanion) and the right side (polycation) syringes by two 2.5 mL transfer pipettes (SARSTEDT, Germany), respectively. After assembling the spray device, the spray distance between the substrate and the spray nozzle was controlled by a ruler (10 cm and 15 cm). The spray pattern started from left to right (as one layer) and then right to left (as the second layer), repeating this spray pattern until both solutions were deposited onto the targeted substrates. The spray-coated samples were then air-dried and sterilized under UV light inside a class II microbiological safety cabinet (Thermo Scientific, USA) for 30 minutes before *in vitro* antimicrobial and cytotoxicity experiments.

### Spray applicator design, development, and testing

#### Spray design/development

The design and development phase was proceeded in two phases: mechanism design and ergonomic optimization. Mechanical design focused on developing the fine mist spray outcome by a simple, easy-to-use mechanism. Ecodesign principles and the McKinsey Design Index informed the ergonomic design process to enhance sustainability. A market study on existing fine-mist delivery systems was carried out to collect the design ideas at an early stage of the design phase.

#### Spray testing

Spray coating experiments on substrates for *in vitro* tests and on mice for *in vivo* tests were conducted by controlling different spray coating parameters: the number of spray passages (as in layer numbers), the spray distance between the spray nozzle and the substrates. The change of polymer concentrations was investigated in *in vitro* tests to find the optimal coating formulation, and then the optimal coating formulation was validated in *in vivo* tests.

### *In silico* study of interactions in the PAR30/HA complex

#### Molecular dynamics simulation

Molecular dynamics (MD) simulation was used to study the behavior of the PAR30/HA complex in an aqueous solution. The molecular system included 1 100 kDa linear HA molecule (containing 264 N-acetyl-D-glucosamine residues and 263 D-glucuronic acid residues) and 17 PAR30 molecules. We used the doGlycans tools ^74^ to build the input structure of the HA and generate the topology using the OPLS-compatible GLYCAM06 forcefield ^75^. The structure of the PAR30 was built with AmberTools23 (Case DA, 2023), and the topology was generated using the OPLS-AA forcefield ^76^. We defined the triclinic box around the PAR30/HA complex, which was subsequently solvated with water molecules and filled with sodium and chlorine ions to reach the physiological concentration (150mM) and neutralize the system’s charge. All simulations were conducted under Periodic boundary conditions and with the TIP3P water model; all h-bonds were constrained with the Linear Constraint Solver (LINCS) algorithm. The V-rescale thermostat was used to maintain the system temperature at 300 K, and the C-rescale barostat was used to control the pressure at 1 bar. MD was integrated using a leapfrog integrator with a time step of 2 fs, and snapshots were saved for every picosecond. Before the production simulation, the molecular system was minimized and then equilibrated with restraints on the solute heavy atoms under NVT and NPT ensembles for 0.5 and 1 ns, respectively. Completion of equilibration was estimated by the convergence of temperature, pressure, and density parameters. The production simulation with no restraints was run for 100ns. All MD simulations were performed using the GROMACS 2024 ^77^.

#### Analysis of interactions

Frames from the MD production run were analyzed for the presence of specific non-covalent interactions between PAR30, HA, and ions. For the interaction detection, we used the *arpeggio* Python package ^78^. The analyzed interactions included polar interactions, hydrogen bonds, hydrophobic interactions, ionic interactions (salt bridges), and others available in the *arpeggio* package.

### Coating characterization

#### Focused Ion Beam cross-sectional analysis

*FIB* cross-sectional analysis was performed to evaluate the coating thickness, uniformity, and elemental distribution on medical-grade glass and Ti40 substrates at a working distance of 4 mm and a tilt of 52°. A FEI HELIOS NanoLab 600i Multi-Beam FIB-SEM system (FEG-SEM and Ga ion columns) equipped with an Oxford Instruments SDD EDS detector and AZtec software was used for elemental mapping. Prior to analysis, a ∼20 nm carbon layer was deposited by evaporation (LEICA EM ACE600) to enhance surface conductivity. Cross-sectional imaging was carried out in both secondary electron (SE) and backscattered electron (BSE) modes. Before milling, a protective platinum (Pt) layer (∼20 µm length × 2 µm width × 2 µm thickness) was deposited in situ to minimize ion-induced damage. Elemental characterization was conducted at 10 kV for uncoated substrates and 5 kV for coated samples. The cross-sections clearly revealed distinct coating layers, including the Pt protection layer, the polymer coating, and the underlying Ti40 alloy, and glass substrates.

#### Raman spectroscopy

Confocal Raman spectroscopy (HORIBA Jobin Yvon, France) was employed for chemical mapping of the 10P5H coatings on glass and Ti40 substrates. Samples were excited with a 532 nm laser under a ×100 objective, providing sub-micron lateral resolution. Spectra were collected over 200–4000 cm ¹, calibrated against a silicon standard, baseline-corrected, and analyzed using LabSpec 5 software to determine the composition of the coatings and substrates.

### *In vitro* antimicrobial test

#### Bacterial strains

Three bacterial strains were selected: *E. coli* ATCC 8739, *S. aureus* ATCC 25923, and *P. aeruginosa* ATCC 9027. Bacteria were stored using ceramic bead cryopreservation and revived by rolling a single bead onto Mueller-Hinton (MH) agar (ThermoFisher Scientific), followed by incubation at 37 °C for 24 h in a CO incubator (ICO, Memmert).

#### Antimicrobial testing

Preculture made with an isolated colony was transferred into 5 mL MH broth and incubated overnight at 37 °C with shaking (Titramax 101, Heidolph). Glass slides were sterilized under UV light for 30 min. To simulate worst-case contamination, the inoculum was adjusted to 10 CFU/cm², exceeding the ISO 22196 “Plastics - Measurement of antibacterial activity on plastic surfaces” of 10 CFU/cm². Samples are incubated with this inoculum for 24h at 37°C with shaking. Bacterial growth was assessed via optical density (OD) using a MultiSkan SkyHigh plate reader (Thermo Scientific). Supernatants were transferred in triplicate to a 96-well plate for absorbance measurement. For viable count analysis by CFU counting, slides were washed thrice with phosphate-buffered saline (PBS), transferred to Falcon tubes containing PBS, vortexed (MS 3 digital, IKA) for 30 s, sonicated for 1 min (XUB12, Grant Instruments), and vortexed again for 30 s. Serial dilutions were spread on MH agar and incubated at 37 °C for 24 h.

#### Sterilisation impact assessment

Solutions were autoclaved using the autoclave DE200 from Systec, at 121.4°C during 20min plateau. Then, solutions are tested as described above using *S. aureus* ATCC 25923, and *P. aeruginosa* ATCC 9027. For this section, only the optical density was measured.

### Drop plate method

#### Plate Coating

Sterile 9 cm Petri dishes (65 cm²) containing Mueller–Hinton (MH) agar were coated using the spraying device. Syringes were filled with 2.5 mL of (i) Tris–NaCl buffer (10–150 mM, 0.2 µm filtered; control), (ii) PAR30 (10 mg/mL), (iii) HA144 (5 mg/mL), and (iv) antibiotics (Tetracycline 100µg/mL and Cefotaxime 1µg/mL), and sprayed from 10 cm. One spray was used to coat either one (65 cm²) or two dishes (130 cm²). Plates were dried under a sterile laminar flow hood (∼1-4 h) and used immediately or stored at 4 °C until inoculation. *P. aeruginosa ATCC 27853*, *S. aureus ATCC 25923, and E. coli ATCC 25922* overnight cultures were serially diluted 10-fold in sterile PBS. Aliquots (5 µL) of undiluted cultures and selected dilutions were spotted in triplicate onto coated and uncoated MH agar plates and incubated overnight at 37 °C.

#### Colony Enumeration

Colonies were counted after incubation. With a deposited volume of 5 µL, the limit of detection was 200 CFU/mL. Counts from spots containing 10–40 colonies were used to calculate mean values and determine microbial concentrations (CFU/mL) based on the dilution factor.

#### Ageing assay

Tests were conducted on 18-month-old PAR30 and HA solutions. Both polymer solutions were prepared in an ISO 7 cleanroom under an ISO 5 laminar air flow. The solutions were stored between 2°C and 8°C in glass bottles or in syringes being prefilled in an ISO 7 cleanroom. Antimicrobial activity was assessed using the drop plate method explained above on exponential *S. aureus* precultures.

#### *In vitro* cytotoxicity test

The mouse embryonic fibroblast cell line Balb/3T3 clone A31 (ATCC CCL163), known for its high sensitivity in cytotoxicity assays, was used as the test model.

They were cultivated in Dulbecco’s Modified Eagle’s Medium (DMEM) (Dutscher) as a base, adding 10% Fetal Bovine Serum (FBS) (Dutscher) and 1% of antibiotics penicillin/streptomycin (Dutscher). Cell culture medium and 20% Dimethyl sulfoxide (DMSO) (Fisher scientific) were used as positive and negative controls respectively. Coated glass slides were prepared using the spray-coating protocol and were placed in contact with Balb/3T3 cell monolayers at 180°000 cells/well and incubated for 24 h under standard conditions (37 °C, 5% CO). The Alamar Blue test was performed using resazurin powder (Thermo Scientific) diluted in PBS to 75 mM, corresponding to a 1000X concentration. After a 24h incubation the medium was removed and replaced by the resazurin solution diluted 1/10 into culture medium. The plate was incubated in the dark at 37°C, 5% CO2 for 2 h. Finally, supernatants from the 6-well plate were transferred in triplicate into a black 96-well plate and fluorescence was measured using Varioskan Lux (Thermo Scientific) at 530_ex_/590_em_ nm to assess the cell viability.

### Skin sensitization and irritation

#### Sample preparation

Poly-L-Arginine and hyaluronic acid were solved in DMEM (low glucose) supplemented with 10% fetal bovine serum and 1% L-glutamine, in medium containing 4% DMSO at a stock concentration of each 400 µg/mL.

#### Keratinosens Assay

KeratinoSens cells were maintained in supplemented DMEM (low glucose) at 37 °C and 5% CO and used between passages 10 and 25. Cells were seeded at 10,000 cells per well in 96-well plates and exposed after 24 h to serial dilutions of samples Poly-L-Arginine and hyaluronic acid (final concentration, 0.196–400 µg/mL). Solvent control (4% DMSO) as negative control and additionally ethylene glycol dimethacrylate (EGDMA) as a positive control were included in this study. After 48 h, cell viability and luciferase induction were measured using alamarBlue and One-Glo, respectively, on a Synergy H1 microplate reader.

### In vivo wound model

#### Animals

Seven-week-old female BALB/c mice (Charles River Laboratories), weighing between 15 and 21 g, were used for the study. Mice were housed under specific pathogen-free conditions, in standard cages, and maintained on a 12-hour light/dark cycle with ad libitum access to food and water. All experimental procedures were conducted in accordance with institutional guidelines and approved animal care protocols (Institut Pasteur AC466_23.207).

#### Wound Creation, spray application, and infection

Each mouse received two standardized full-thickness skin wounds, each 5 mm in diameter, created on the dorsal surface using a sterile skin punch biopsy tool. Wounds assigned to the experimental group were immediately treated with the antimicrobial spray coating (the same “spray coating protocol” mentioned above). Control wounds remained untreated. Following the spray application, a bacterial inoculum of *S. aureus* Xen31 (MRSA strain, ATCC 33591) was deposited directly onto the wound surface at a concentration of 1 × 10 CFU per wound. All wounds were immediately covered with a sterile gauze compress after bacterial inoculation.

#### Bacterial load assessment and imaging

To monitor bacterial colonization and biofilm formation, bioluminescent imaging was performed using the IVIS Spectrum imaging system (Caliper Life Sciences, Hopkinton, MA, USA). Mice were anesthetized with isoflurane during imaging sessions. Images were captured immediately after bacterial inoculation and spray application, and subsequently every other day up to day 15 or until the study endpoint. Bioluminescence was quantified as average radiance (photons/sec/cm²/steradian) using a standardized region of interest (ROI) across all images. In addition, at selected time points (notably at 48 hours post-infection), bacterial burden was evaluated by recovering both wound tissue and dressing materials. Gauzes were removed aseptically, and wound sites were excised. Tissue samples were homogenized in sterile PBS, serially diluted, and plated onto nutrient agar for colony-forming unit (CFU) enumeration.

#### Statistical analysis

Bioluminescent signal intensities and CFU counts were analyzed using GraphPad Prism software. For comparisons between groups, two-way analysis of variance (ANOVA) followed by multiple comparisons was applied for imaging data, and unpaired Student’s t-tests were used for CFU data. Results are presented as mean values ± standard error of the mean (SEM). Statistical significance was considered at p < 0.05.

### Nociceptive and motor behavioral assays in a post-surgical pain model

#### Animals

Experiments were performed at the Chronobiotron facility (UMS3415; approval no. D67-2018-38) using male C57BL/6J mice (Charles River, France) aged six weeks at the start of the study. Animals were group-housed (four per cage) under a 12 h light/dark cycle with food and water *ad libitum*, in cages enriched with nesting material and wooden gnawing bars. Body weight was monitored regularly, and experiments commenced after a 1-week habituation period. All procedures were approved by the Regional Ethics Committee for Animal Experimentation of Strasbourg (CREMEAS CEEA 035).

#### Treatment procedures

Mice paws were sprayed with a combination of hyaluronic acid (HA HTL, 5 mg/mL) and polyarginine (PAR30, 10 mg/mL) solution or with saline (NaCl 0.9%). The spray device was equipped with two 2.5 mL syringes, filled either with 1 mL each of HA HTL and PAR30 solutions or with 1 mL of saline solution in each syringe for the control group.

For the paw incision groups, four spray applications were performed: the first application was on the muscle after the paw incision, the second on the skin immediately after suturing (T0), and the third and fourth on the skin 2 hours (T2) and 24 hours (T24) after the second application, respectively. In the no-incision groups, only three skin spray applications were performed at T0, T2, and T24.

#### Paw incision procedure

Mice were anesthetized by isoflurane inhalation (5% for induction and 2.5% for maintenance). The plantar surface of the right hind paw was prepared under sterile conditions, and a midline incision was made using a No. 11 blade, starting 0.5 cm from the heel. The wound was closed with 5-0 nylon mattress sutures.

#### Scoring

Behavioral scoring was performed immediately after the third (T2) and fourth (T24) spray applications. Mice were placed in transparent Plexiglas boxes (7 cm × 9 cm × 7 cm) on a raised metal grid and licking and shaking behaviors were recorded over a 3-minute period.

#### Von Frey test

The mechanical nociceptive threshold was evaluated using the Von Frey filament test. Mice were placed in transparent Plexiglas boxes (7 cm × 9 cm × 7 cm) on a raised metal grid and allowed to habituate for 15 minutes. Calibrated nylon filaments (0.4–8 g) were applied to the plantar surface of each hind paw in ascending order of force. The mechanical threshold was defined as the lowest filament force eliciting at least three withdrawals in five successive applications ^79^. Results were expressed in grams.

#### Dry ice test

Thermal nociceptive sensitivity to cold was assessed using the Dry Ice test. Mice were placed in transparent Plexiglas boxes (7 cm × 9 cm × 7 cm) on a glass surface and allowed a 10-minute habituation period. A syringe containing dry ice (0.5 cm diameter) was placed under the plantar surface of each hind paw. Paw withdrawal latencies were measured, and three measurements were averaged for each paw. A 20-second cut-off was set to prevent tissue damage.

#### Hargreaves test

Thermal nociceptive sensitivity to heat was evaluated using the Hargreaves method. Mice were placed in transparent Plexiglas boxes (7 cm × 9 cm × 7 cm) on a glass surface and habituated for 10 minutes. A fiber-optic infrared heat source was applied under the plantar surface of each hind paw. Paw withdrawal latencies were measured, and three measurements were averaged for each paw. A 20-second automatic cut-off was implemented to prevent skin damage.

#### Beam walking test

Motor coordination was assessed at days 6 and 7 post-surgery using a beam walking task, in which mice traversed an elevated horizontal bar (80 cm) of decreasing diameter (2.0 cm on day 1; 1.2 cm on day 2) to reach a goal box. Performance was evaluated by the latency to traverse and the number of paw slips across increasing distances (5–100 cm).

#### Statistics

Graphs were created using GraphPad Prism 9 software. Data were expressed as group means ± standard error of the mean (SEM). Statistical analysis was performed using two-way analysis of variance (ANOVA) followed by Tukey’s multiple comparisons test using GraphPad Prism.

### SSbD

#### The SSbD Assessment

A simplified SSbD approach was followed, as the biopolymer solution was in the early stages of innovation, considering.^21,80^. The composition of the solution can be seen in Supplementary Table S9. The SSbD focuses only on the biopolymer solution itself, and the assessment covers the hazard classification, occupational health and safety during processing, human and environmental risks during the use phase, and sustainability aspects of production of the solution, while it excludes the socio-economical assessment, end-of-life considerations, and environmental impacts from transportation and the applicator.

#### Hazard classification

The results from *in vitro* and *in vivo* tests, together with the most relevant hazard category is skin sensitization, as the solution is directly applied to the wound. OECD #442D: In Vitro Skin Sensitization tests were applied ^37^, and concentrations between 0.19 – 400 µg/mL were tested.

#### Occupational health and safety during processing

The StoffenManager tool (https://stoffenmanager.com), a software designed to support safe and healthy working conditions when handling hazardous substances, was used. It serves as a self-assessment tool developed by the Netherlands Labour Authority to help organizations comply with legal requirements. The considerations, together with the assumptions made during this step, are presented in the Supplementary Table S10.

#### Risk assessment; human and environmental health

For the exposure concentrations, worst-case scenarios were considered. The human exposure scenario is as follows: the biopolymer solution is applied once per day to the patient’s wound in a hospital room, and it is assumed that 100% of the solution is inhaled. Environmental exposure was calculated based on the sales projection for 2030, assuming a 100% release scenario. For the human hazard assessment, Derived No-Effect Level (DNEL) values were derived from the literature. The risk characterization ratios (RCR) for human health are calculated by using “Exposure concentration” divide “Hazard threshold (DNEL)”. If the RCR is below 1, no risk is expected, whereas an RCR equal to or greater than 1 indicates a potential risk. For the environmental compartments, the risk analysis was based on the direct comparison between the exposure concentrations and ecotoxicity data (NOVA Deliverable Report 6.2 ^81^), without the application of additional factors as the comparison of exposure and hazard data alone was sufficient to conclude a low likelihood of adverse ecological effects due to the substantial margin between the two. The data used, together with the assumptions made, can be found in Supplementary Table S5.

#### Sustainability assessment

Life cycle assessment (LCA) was conducted using the Activity Browser, which is an open-source software ^82^. Based on the described system, the inventory to produce the biopolymer solution was collected, and the ecoinvent database v3.11 (https://ecoinvent.org/) was used to model the processes. Ecoinvent contains over 25,000 datasets representing human activities and processes across various sectors and is widely used in LCA. When the related chemical was unavailable in the database, relevant ingredients were used as proxies. Two different sets of proxies were used for HA and PAR, as presented in Supplementary Table S6, in order to minimize the possibility of false-positive direct impacts linked to a single proxy. The Environmental Footprint v3.1 method was used for the impact assessment. The functional unit was defined as "keeping 65 cm^2^ wound antimicrobial for 24 hours". A simplified LCA approach was followed, in which only the production and processing of the solution were considered, while other processes (e.g. transportation, applicator) were excluded. A comparative analysis was done with polyhexanide-betaine solution as the main alternative to the biopolymer solution. Additional considerations originating from the biocide-specific SSbD framework were also considered, as shown in Supplementary Table S8. The sources of the data used for the SSbD assessment can be found in Supplementary Table S11.

## Data Availability

Available upon request

## Acknowledgements

We would like to thank the entire SPARTHA Medical team for their contribution to this work. We thank Tammi Daoudi and Fabrice Thiebaud for their help in designing the first spray device used in this study. This work benefits from a National Research Agency (ANR) aid under the investment program for the future integrated to France 2030 bearing the reference ANR-23RHUS-0004. The EIC Accelerator project SPARTHACUS, the Horizon Europe project NOVA, and BPI through the Aide DeepTech programme.

## Ethics declarations

Some contributors of this work are employees of SPARTHA Medical.

