## Supplemental Figures and Tables for "Invisible shield: Sprayable supramolecular antimicrobial microscale films for preventing wound and medical device infections"

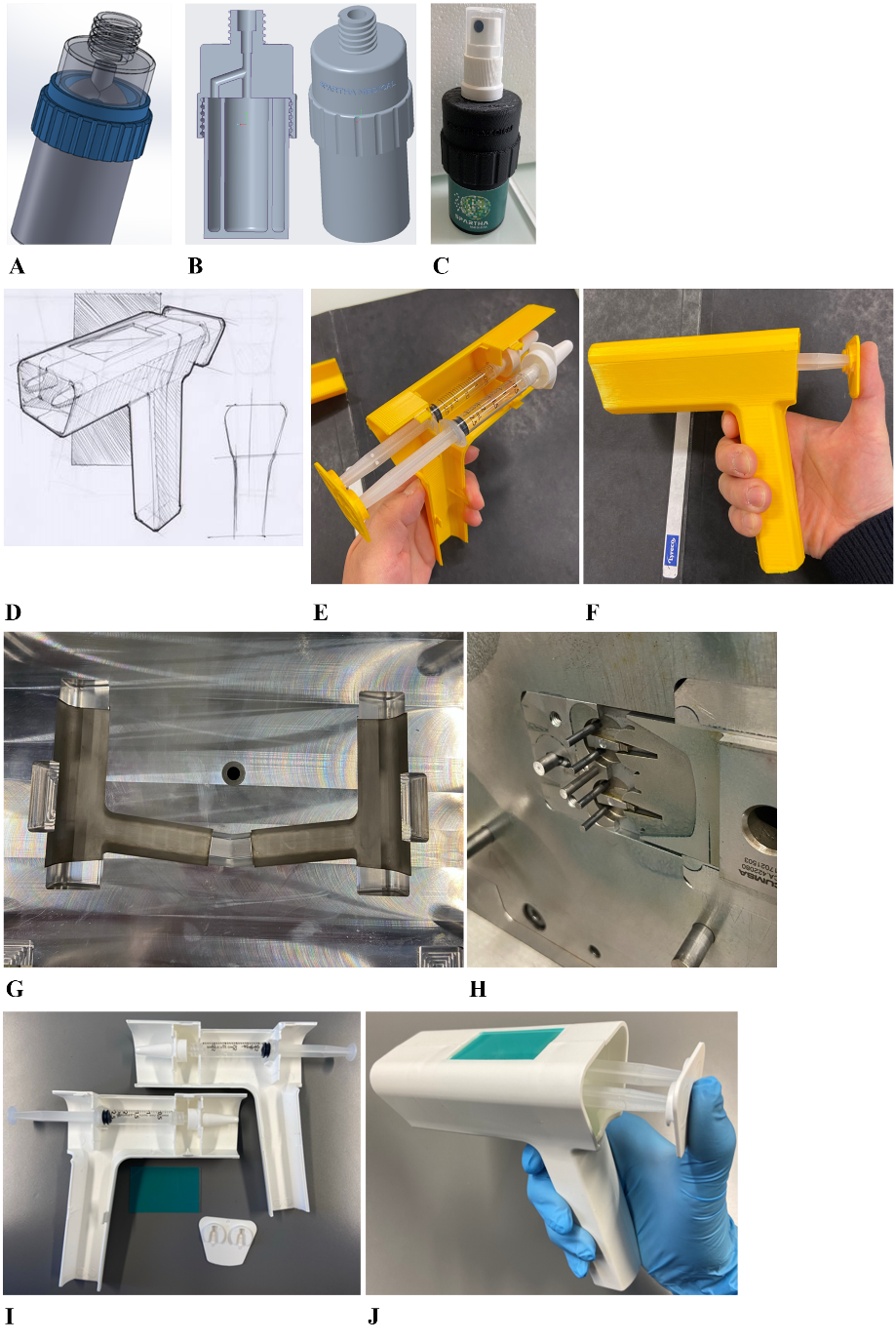

**Supplementary Figure S1.** **Development of a portable biopolymer delivery system (A)** 3D model of the double-output spray bottle – First generation. **(B)** Updated 3D model of the spray bottle. **(C)** 3D printed prototype of the spray bottle. **(D)** Updated design of the syringe spray system. **(E)** Disassembled 3D printed handhold piece. **(F)** Assembled 3D printed handhold piece. **(G)** Injection mold for the handhold piece. **(H)** Injection mold for the back pusher. **(I)** Disassembled final product. **(J)** Assembled final product.

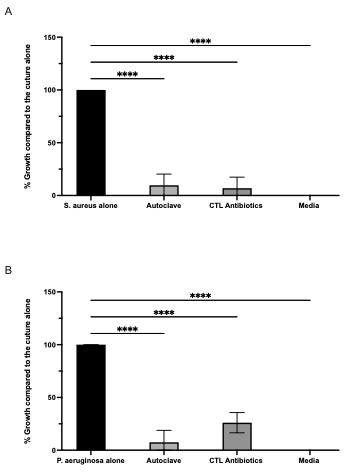

**Supplementary Figure S2.** **Antibacterial activity of autoclaved PAR30 and HA solutions**. PAR30 (10 mg/mL) and HA (5 mg/mL) solutions were autoclaved at 121.4°C for 20 min plateau. Antibacterial activity against *S. aureus* **(A)** and *P. aeruginosa* **(B)** was evaluated after 24 h. Bacterial growth is expressed as percentage relative to the bacteria-only control. Data represent mean ± SD (n = 3). Statistical analysis was performed using one-way ANOVA, with significance indicated as *****p* < 0.0001.

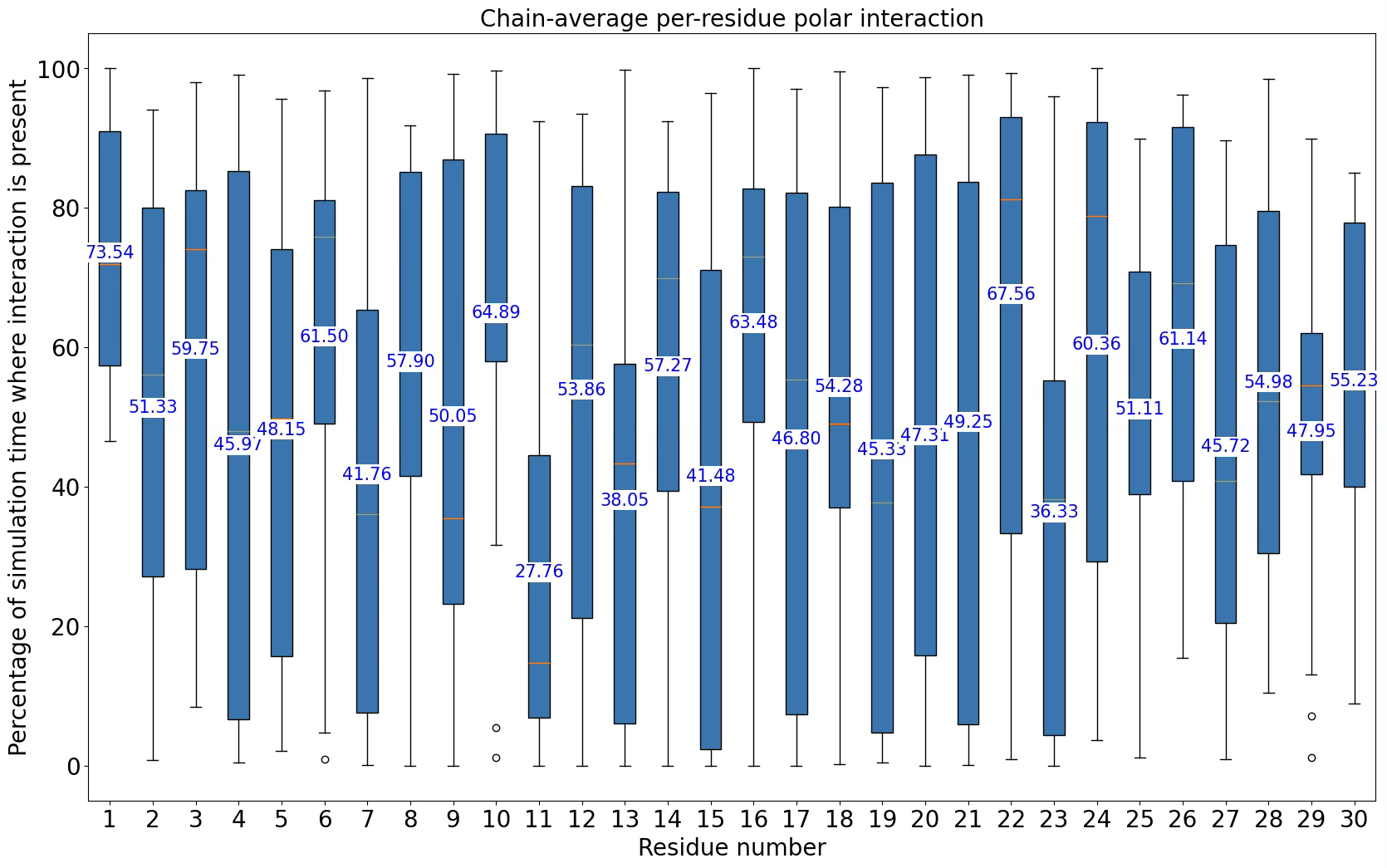

**Supplementary Figure S3**. **Molecular dynamics (MD) simulations.** Polar interactions (also includes ionic/charged interactions) between arginine residues at different positions in PAR30 chains with HA. The values on the boxplots are means calculated across 17 PAR30 chains present in the simulation.

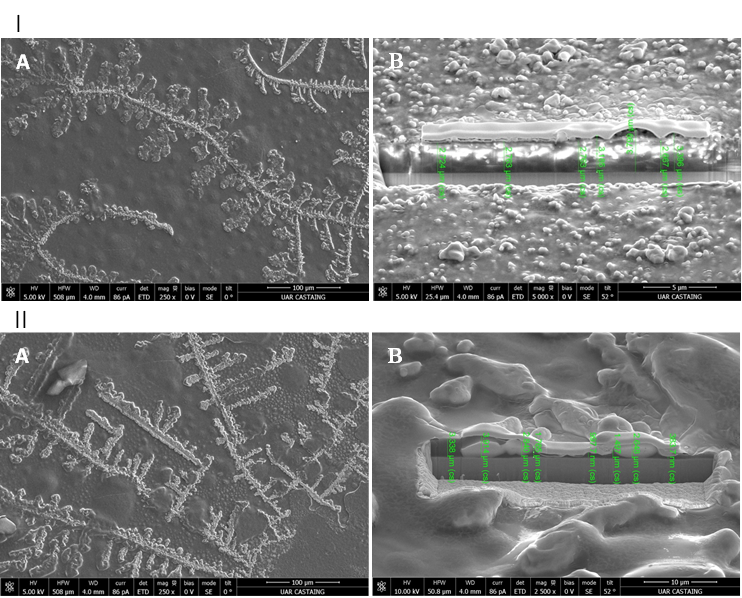

**Supplementary Figure S4. FIB–SEM images of 10P5H-coated samples.** Panels show **(I)** glass and **(II)** Ti40 substrates. Images **(A)** Surface morphologies (mag. 250×), and images **(B)** cross-sectional views of the coatings, illustrating coating continuity and thickness.

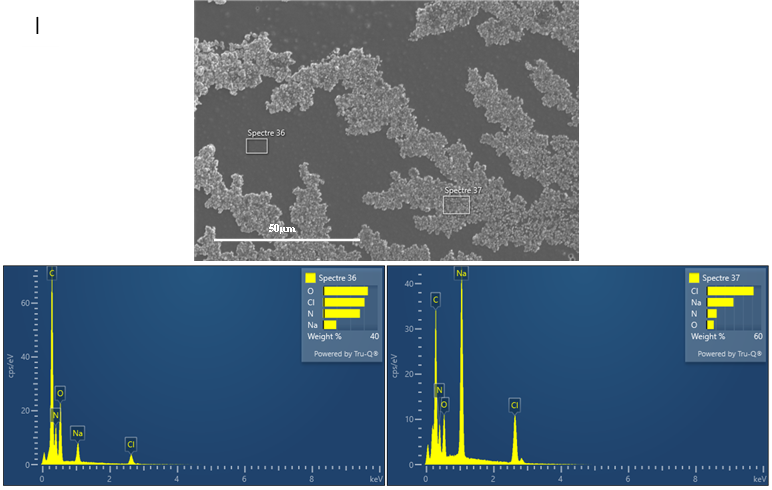

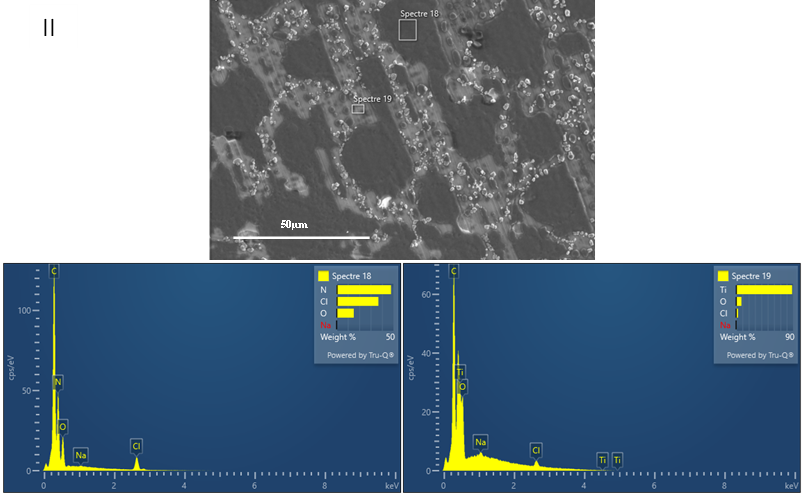

**Supplementary Figure S5. FIB–SEM/EDS analysis of 10P5H-coated samples.** Panels show results for **(I)** glass (spectra 36 and 37) and **(II)** Ti40 (spectra 18 and 19) substrates. EDS analyses were performed at an accelerating voltage of 5 kV using a probe current of 1.4 nA for glass samples and 2.8 nA for Ti40 samples. Elemental compositions (at.%) are indicated for each spectrum. Standard background subtraction was applied during data processing.

**Supplementary Table S1.** Design specifications for the portable antimicrobial spray device.

| Design elements | Requirements description | Validation method |
| --- | --- | --- |
| Coating efficacy | Biocompatible | *In vitro* cytotoxicity test |
|  | Antimicrobial ability (at least 3 log reduction) | *In vitro* antimicrobial test  *In vivo* antimicrobial test |
|  | Have a barrier presence | Confocal microscope images on substrates, FTIR analysis on substrates |
|  | No delay in wound healing | *In vivo* antimicrobial test |
|  | Pain reduction | *In vivo* pain study |
| Functionality | Simultaneous spray of two biopolymers | Usability test with visual check during usage |
|  | Mixture on the substrate | Usability test with coating images as validation |
|  | Fine mist outcome | Usability test with visual check during usage |
|  | Spray distance control and validation | Usability test with validated antimicrobial results |
|  | Volume control and validation | Usability test with validated antimicrobial results |
| Ergonomic | Easy to press | Usability test on user feedback |
|  | Easy to hold | Usability test on user feedback |
|  | Easy to discard | Usability test on user feedback |

**Supplementary Table S2.** Atomic percentages of elements in 10P5H-coated and uncoated glass and Ti40 substrates, determined by FIB–EDS.

| Sample | Spectrum | N | O | Na | Cl | Ti | Mg | Si | k | Ca | Sn |
| --- | --- | --- | --- | --- | --- | --- | --- | --- | --- | --- | --- |
| Coated Glass | Spectre 36 | 35 | 39 | 8 | 18 | – | – | – | – | – | – |
|  | Spectre 37 | 19 | 12 | 30 | 39 | – | – | – | – | – | – |
|  | Average | 27 | 25.5 | 19 | 28.5 | – | – | – | – | – | – |
| Uncoated Glass | De carte | – | 61 | 7 | – | – | 2 | 24 | 0.3 | 3.8 | 1.5 |
| Coated Ti40 | Spectre 18 | 50 | 19 | – | 17 | 14 | – | – | – | – | – |
|  | Spectre 19 | 4 | 21 | – | 4 | 71 | – | – | – | – | – |
|  | Average | 27 | 20 | – | 10.5 | 42.5 | – | – | – | – | – |
| Uncoated Ti40 | Spectre 1 | – | 11 | – | – | 89 | – | – | – | – | – |

**Supplementary Table S3. Evaluation of skin sensitization and irritation potential of Poly-L-arginine and hyaluronic acid according to the OECD TG 442D.** *EC1.5: Concentration inducing a 1.5-fold luciferase response; IC30 and IC50, concentration causing 30% and 50% reduction in cell viability, respectively; n.d., not determined.*

| **Sample** | **EC_1.5_ (µg/mL)** | **IC_30_ (µg/mL)** | **IC_50_ (µg/mL)** | **Interpretation** |
| --- | --- | --- | --- | --- |
| Poly-L-arginine | n.d. | 6.41 | 33.52 | Potential irritant |
| Hyaluronic acid | >400 | >400 | >400 | Not sensitizer nor irritant |

**Supplementary Table S4. Results of the risk Assessment conducted using Stoffenmanager®.** *The results show potential hazards to skin, eyes, and via inhalation, along with recommendation control measures and risk prioritization.*

| **Risk assessment - skin** | | | |
| --- | --- | --- | --- |
| **Type** | **Hazard class** | **Exposure class** | **Risk priority (/3)** |
| Skin local (risk skin contact with substance) | - | 3 - average | Third priority |
| Skin uptake (risk uptake of substances through the skin) | - | 2 - low | Third priority |
| **Risk assessment - eyes** | | | |
| **Hazard class** | **Recommended eye protection** | | **Remark** |
| B - average | Adequate eye protection is highly recommended: wear safety glasses with side protection | | Depending on the task, the recommended eye protection is not always needed |
| **Risk assessment - inhalation** | | | |
| **Hazard class** | **Exposure class** | | **Risk priority** |
| - | 1 - Low | | Third priority |

**Supplementary Table S5. Risk characterization approach**

Exposure scenario human health

*Assumptions:*

Density of the solution ≈ 1 g/mL.

*Approach:*

Concentration per application: HA: 0.013 mL, 13 mg; PAR per application: 0.025 mL, 25 mg

Concentration in the room: HA: 0.37 mg/m^3^; PAR: 0.7 mg/m^3^

Inhalation rate: 0.36 m³/h (Ref) (exposure: 24 hours): 8.64 m³

Inhaled dose: HA: 3.2 mg (8.64 m³ x 0.37 mg/m^3^); PAR: 6 mg (8.64 m³ x 0.7 mg/m^3^)

Total cumulative dose in 5 days: HA: 16 mg/m^3^; PAR: 30 mg/m^3^

Derived No-Effect Level (DNEL) values

*PAR:*

DNEL values for L-Arginine monohydrochloride were used 668.2 mg/m^3^ (Carl Roth GmbH + Co. KG, 2024), as no

DNEL value cannot be found for Poly(L-arginine hydrochloride).

*HA:*

HA has been used in medical sector for treatment purposes including airway diseases (Carro and Martínez-García, 2020), and is considered as non-toxic (Tiwari and Bahadur, 2019). Since no DNEL values were found for HA, medical data from the literature was used. Macchi et al. (2013) used intranasal 9 mg nebulised sodium hyaluronate in isotonic saline twice per day in patients, and no adverse effects were reported. It is assumed that 18 mg of hyaluronic acid is safe (The weight of hyaluronic acid and sodium hyaluronate assumed to be same, and 1m3 air is considered).

Risk characterization ratios (RCR):

RCR PAR: 30 mg.m^-3^/ 668 mg.m^-3^ : 0.05

RCR HA: 16 mg.m^-3^/18 mg.m^-3^: 0.88

Exposure scenario environment

*Assumptions:*

100% release, no degradation.

Density of the solution ≈ 1 g/mL.

*Sales projection:*

138,000 pieces in 2030; 690 L/year (138,000 x 5 mL/piece)

Volume of PAR and HA is 3.4 and 1.7 L/year, respectively.

The substance release: PAR: 0.0093 kg/day, HA: 0.0046 kg/day.

Exposure aquatic environment: (calculated considering ECHA, 2016)

PAR: 0.0093/(2000.10): 0.465 mg/L

HA: 0.0046/(2000.10): 0.23 mg/L

Hazard values used (NOVA Deliverable Report 6.2, 2025)

*Tetrahymena thermophila*;

PAR30: LOEC: 10 mg/L,

HA100: HONEC: 200 mg/L

*PNEC:*

*Daphia magna*;

PAR30: LOEC: 40 mg/L,

HA100: HONEC: 40 mg/L

*Porcellio scaber*; PAR30: HONEC: 10 g/kg, HA100: HONEC: 10 g/kg

Risk analysis:

Direct comparison of exposure concentrations and ecotoxicity data indicates a low likelihood of environmental risk on aquatic and terrestrial systems.

**Supplementary Table S6. Life Cycle Assessment Results.** *Biopolymer solution performance compared to alternative chemicals. The numbers were not reported on purpose since it is a preliminary assessment to show a general overview.*

| Impact Category | Set 1 | Set 2 |  |  |
| --- | --- | --- | --- | --- |
| Acidification \| accumulated exceedance (AE) |  |  |  | Higher impact |
| Climate change \| global warming potential (GWP100) |  |  |  | Lower impact |
| Ecotoxicity: freshwater \| comparative toxic unit for ecosystems (CTUe) |  |  |  | No change |
| Eutrophication: freshwater \| fraction of nutrients reaching freshwater end compartment (P) |  |  |  |  |
| Eutrophication: marine \| fraction of nutrients reaching marine end compartment (N) |  |  |  |  |
| Eutrophication: terrestrial \| accumulated exceedance (AE) |  |  |  |  |
| Human toxicity: carcinogenic \| comparative toxic unit for human (CTUh) |  |  |  |  |
| Human toxicity: non-carcinogenic \| comparative toxic unit for human (CTUh) |  |  |  |  |
| Ionising radiation: human health \| human exposure efficiency relative to u235 |  |  |  |  |
| Land use \| soil quality index |  |  |  |  |
| Ozone depletion \| ozone depletion potential (ODP) |  |  |  |  |
| Particulate matter formation \| impact on human health |  |  |  |  |
| Photochemical oxidant formation: human health \| tropospheric ozone concentration increase |  |  |  |  |
| Water use \| user deprivation potential (deprivation-weighted water consumption) |  |  |  |  |
| Energy resources: non-renewable \| abiotic depletion potential (ADP): fossil fuels |  |  |  |  |
| Material resources: metals/minerals \| abiotic depletion potential (ADP): elements (ultimate reserves) |  |  |  |  |

**Supplementary Table S7. Life Cycle Inventory.** *As not all the ingredients can be found in the database, their selected alternatives were shown. Different alternatives (e.g. glucose as raw material, and ethanol as the product of fermentation) were used to reduce the dependency on the single process. Tris base was assumed to be neglectable due to low amount used. Background system in the database was considered. European data were preferred, if available.*

| Main ingredient | Set 1 | Set 2 |
| --- | --- | --- |
| Hyaluronic acid | Glucose | Ethanol |
| Polyarginine | Glucose + ammonium sulfate ^1^ | Ethanol |
| HCl | HCl | HCl |
| NaCl | NaCl | NaCl |
| Water (distilled) | Water (deionized) | Water (deionized) |
| Polyhexanide | Hexamethylenediamine  + heat ^2^ | Hexamethylenediamine  + heat ^2^ |
| Betaine | Organic solvent + heat ^3^ | Organic solvent + heat ^3^ |

^1^ as it is another important raw material (Takashi et al. 2004).

^2^ as heat is needed for the synthesis (Wang et al,, 2023).

^3^ as heat is needed for the synthesis (VulcanChem, 2026).

**Supplementary Table S8.** **Additional considerations**

*Approach*

Additional considerations from biocide-specific framework include considering i) the possibility of developing bacterial -resistance, ii) the possibility and severity of accidents during production and use phases, iii) benefit-based functional unit in sustainability assessment, iv) central aspects, efficacy, minimum sufficient concentration, durability and possibility of less frequent application.

*Results*

Bacterial resistance is shown to be unlikely for the biopolymer solution. PAR30 was tested against developing resistance, and bacteria did not develop a resistance to PAR (Kocgozlu et al., 2024). The role of HA in biopolymer solutions is related to support the PAR chain mobility in the context, and thus is irrelevant for this context . By being a biodegradable polymer with simple processing and application, and is only available for professional use and application no accidents or incidents is foreseen. When it comes to the central aspects, the efficacy and minimum sufficient concentration were tested. It is biodegradable, however, based on vivo test results, it can be applied once in every two days with no negative impact on skin, which may result in lower usage of the solution. However, since it is still early innovation phase, applying once a day is preferable as it is common sense compared to the bandaging.

**Table S9. Ingredients of the biopolymer solution assessed for SSbD.**

| PAR polymer solution | Concentration (%) |
| --- | --- |
| Polymer Component: Poly-arginine | 0.99 |
| Tris Base (Tris(hydroxymethyl)aminomethane) | 0.00012 |
| NaCl (Sodium Chloride) | 0.00087 |
| HCl (Hydrochloric Acid) | 0.86 |
| Distilled Water | 98.15 |
| HA polymer solution | **Concentration (%)** |
| Polymer Component: Hyaluronic acid | 0.5 |
| Tris Base (Tris(hydroxymethyl)aminomethane) | 0.000121 |
| NaCl (Sodium Chloride) | 0.000872 |
| HCl (Hydrochloric Acid) | 0.86 |
| Distilled Water | 98.64 |
| The HA and PAR are mixed in percentage close to 50/50 during usage. | |

**Table S10. Considerations for Stoffen Manager Tool.**

The workspace is not segregated with screens/cabins/walls between the worker and the source.

Distance to the chemical/material source in the working environment is equal to or less than one arm's length.

The source is not completely enclosed (e.g. no shield around the source). ²

No respiratory protection is used.

The default value for the vapor pressure as 2300 Pa at 20 ^o^C were used for the solution.

Boiling point of the solution is 100 ^o^C.

**Supplementary Table S11. Data sources sheet**

| Assessment | Data source |
| --- | --- |
| Hazard Classification | Experimental data |
| Occupational health and safety during processing | Primary data  Assumptions |
| Risk assessment | Primary data  Literature data  Assumptions |
| Sustainability assessment | Primary data  Literature data  Assumptions |
